# An epigenetic barrier sets the timing of human neuronal maturation

**DOI:** 10.1101/2022.06.02.490114

**Authors:** Gabriele Ciceri, Hyunwoo Cho, Meghana Kshirsagar, Arianna Baggiolini, Kelly A Aromolaran, Ryan M Walsh, Peter A Goldstein, Richard P. Koche, Christina S Leslie, Lorenz Studer

## Abstract

The pace of human brain development is highly protracted compared to most other species. The maturation of cortical neurons is particularly slow, extending over months to years to reach adult-like functions. Remarkably, such protracted timing is retained in cortical neurons derived from human pluripotent stem cell (hPSC) during *in vitro* differentiation or upon transplantation into the murine brain *in vivo*. Those findings suggest the presence of a cell intrinsic clock that sets the pace of neuronal maturation, though the molecular nature of such a clock has remained elusive.

Here, we identify an epigenetic developmental program which sets the timing of human neuronal maturation. First, we developed a human PSC-based approach to synchronize the birth of cortical neurons *in vitro* which allowed us to define a detailed atlas of progressive morphological, functional, and molecular maturation in human cortical neurons. Interestingly, we observed a slow, temporal unfolding of maturation programs that is limited by the retention of a specific set of epigenetic factors. Loss-of-function studies for several of those factors in cortical neurons enables precocious molecular and physiological maturation. Remarkably, transient inhibition of EZH2, EHMT1/2 or DOT1L, at the progenitor stage primes newly born neurons to rapidly acquire mature properties upon differentiation. Therefore, our findings reveal that the rate at which human neurons mature is set well before neurogenesis through the establishment of an “epigenetic barrier” in progenitor cells. Mechanistically, this barrier acts by holding transcriptional maturation programs in a poised state that gets gradually released during neuronal differentiation to ensure the prolonged timeline characteristic of human cortical neuron maturation.

## Introduction

The development of the Central Nervous System (CNS) follows a coordinated sequence of events in which a myriad of cell identities is specified, differentiated, and assembled into mature functional circuits. While fundamental developmental steps are broadly conserved throughout mammalian evolution, the pace at which development proceeds is very protracted in humans compared to rodents or even to primates ^1–4^. Interestingly, the *in vivo* sequential order, duration and pace of developmental transitions are largely maintained *ex vivo* during *in vitro* Pluripotent Stem Cells (PSC) differentiation^5^. For instance, PSC from various species differentiated towards lineages of the cerebral cortex faithfully recapitulate the sequential generation of neuron subtypes and glia, following a “schedule” that largely matches the species-specific pace of *in vivo* cortical development^6–9^. Species-specific developmental rates ensue also during the maturation of PSC-derived neurons, with more astonishing (∼10-fold) timing differences between mouse and human neurons compared to the 2 to 3-fold difference in the rate of early embryogenesis^8–14^.

Neuronal maturation represents one of the most lengthy cell transitions that spans fetal and postnatal development and can last weeks, months or even years depending on the species^15^. One striking example is the human cerebral cortex where synaptogenesis and maturation of neural circuits into postnatal development^15, 16^. Similarly, the timeline of hPSC-derived cortical neurons follows a human-specific clock that requires several months to reach mature electrophysiological and synaptic function. The challenge to generate adult-like human neurons is shared across distinct hPSC-derived neuron types, including dopaminergic^17^ and sensory^18^ neurons in addition to cortical excitatory and inhibitory neuron lineages^8, 10, 19, 20^.

While the mechanisms that control neural patterning and cell fate specification during brain development are mostly conserved across species and maintained during PSC differentiation *in vitro*, the mechanisms that set the pace of neural development and neuronal maturation remain largely unresolved. Extrinsic environmental signals, such as neuron-glia interactions^21, 22^, network activity^23^ and secreted molecules^24^ can modulate aspects of neuronal morphogenesis, excitability and connectivity. However, several lines of evidence indicate that the progression towards maturity is primarily timed via cell-intrinsic programs. hPSC-derived cortical neurons transplanted into the rapidly maturing mouse neocortex develop adult-like morphologies, dendritic spine function and connectivity only after about 9 months compared to 4 weeks required for transplanted cortical neurons derived from mouse PSC^10, 25^. Furthermore, grafting of hPSC-derived cortical neurons into mouse models of cortical stroke or midbrain dopaminergic neurons into Parkinsonian rats, required each > 5 months to induce behavioral and functional recovery^17, 26^. Those data indicate that transplanted human neurons follow a species-specific intrinsic timing of maturation *in vivo* rather than adopting the pace of the host species. The protracted maturation of human neurons poses a major challenge for studying neurological and psychiatric disorders that typically manifest as alterations in the activity of postnatal neural networks^27^. Thus, understanding the mechanisms that define and drive the timing of human neuronal maturation is critical to access the full potential of hPSC technologies in modeling and treating brain disorders.

## Results

### A hPSC-based platform to study human neuronal maturation in a dish

A major challenge for the application of stem cell-based models to study human neuronal maturation is the poor synchronization and cell heterogeneity during differentiation. For most differentiation strategies, multiple neuronal lineages coexist with precursor cells which in turn constantly generate newly born neurons that each differentiate at their own pace. Here, we have developed a novel hPSC-based platform that generates homogeneous populations of cortical neurons in a temporally synchronized manner to dissect human neuronal maturation states and their phenotypic complexity in long-term studies (Fig.1a, Extended data Fig.1a). We induced CNS neuroectoderm from hPSC by combined inhibition of TGFβ/Activin/Nodal and BMP signaling pathways (i.e. dual-SMAD inhibition^18^) and optimized cortical patterning by inhibition of WNT signaling^19^ (Extended data Fig.1a). These conditions efficiently coax hPSC, expressing canonical pluripotency markers *OCT4* and *NANOG,* into neural precursor cells (NPC) that express the cortical specific progenitor cell markers *FOXG1*, *PAX6*, *EMX2* and *FEZF2,* among others, by day (d) 10 of differentiation (Extended data Fig.1b). Differentiation into cortical NPC is further shown by robust stage-dependent changes in chromatin accessibility at pluripotency *vs.* forebrain-specific genomic loci (Extended data Fig.1c). Our platform gives rise to a nearly pure, homogeneous population of cortical NPC by d20 (Extended data Fig.1d) that can be triggered towards synchronous neurogenesis based on optimized replating density of cells upon passaging and treatment with the Notch pathway inhibitor DAPT (Extended data Fig.1a, Extended data Fig.2a). By d25 of differentiation, nearly all Ki67^+^ NPC have exited the cell cycle and turned into roughly isochronic MAP2^+^ post-mitotic neurons that are born within only few days apart (Extended data Fig.2b, c), as confirmed by birth-dating analysis with EdU labelling (Extended data Fig.2d, e). Under such conditions, synchronized neurons can be maintained for > 100 days of differentiation with no additional neurogenesis taking place after d25 (Fig.1c, Extended data Fig.2f).

**Fig.1:**
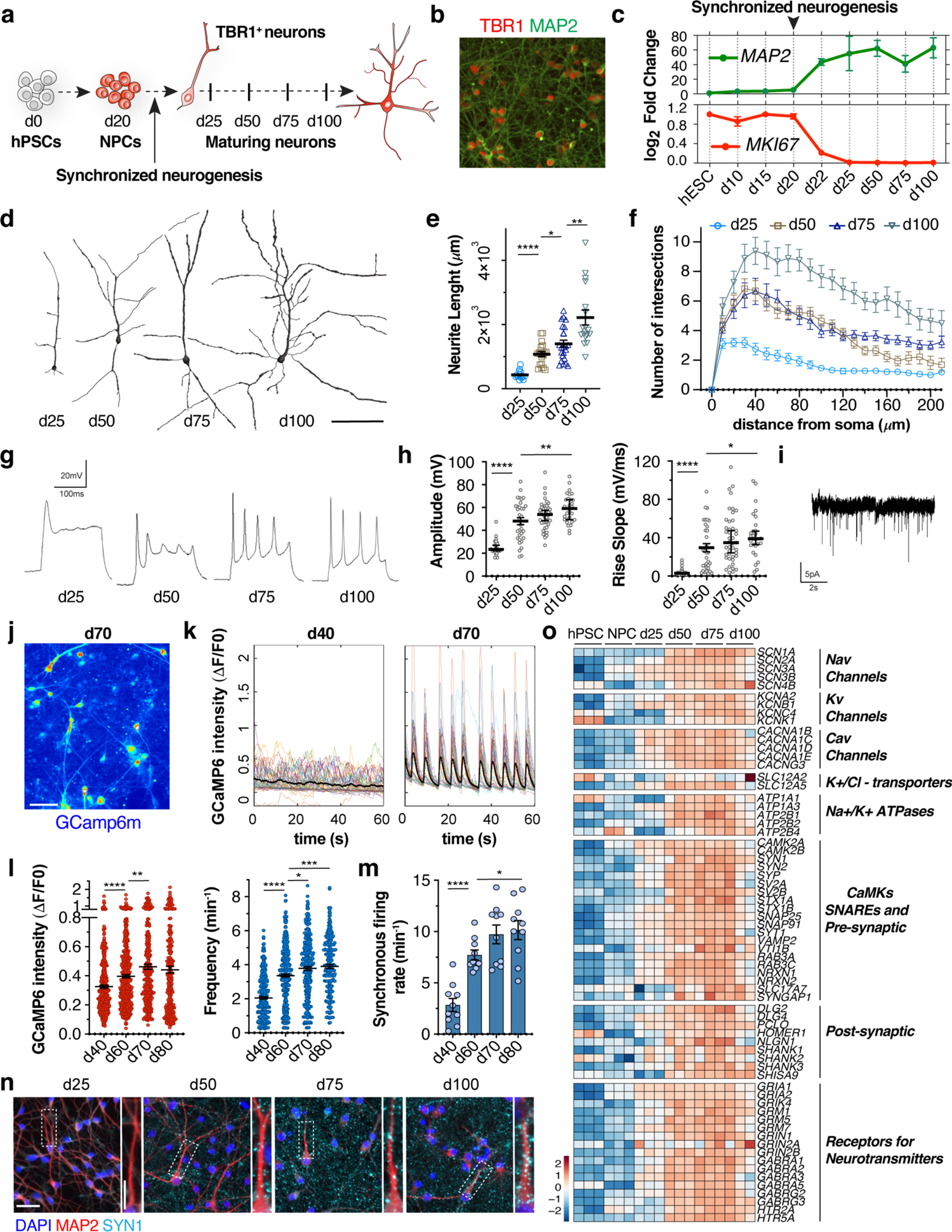
Morphological and functional maturation of synchronized cortical neurons derived from hPSC. **a**, Schematic of the experimental paradigm. **b**, Representative images of neurons stained with antibodies against TBR1. **c**, qRT-PCR expression of *MKI67* and *MAP2* throughout the differentiation (n = 2 independent experiments). Neurons generated under synchronized conditions can be maintained for 100 days *in vitro* without new proliferative events. **d-f**, Representative reconstructions of neuronal morphology (d) and quantification of neurites length (d) and complexity (Sholl analysis, f) during maturation (n = minimum 15 neurons per time point from 2 independent experiments). **g,** Representative traces of evoked action potentials. **h**, Electrophysiological measurements of action potential amplitude and rise slope of cortical neurons over time (n = 25-43 neurons per time point from 10 independent experiments). **i**, Representative traces of mEPSCs at d75. **j,** Representative maximal intensity projection of time-lapse Ca^2+^ imaging at d70. **k,** Representative traces of normalized GCamp6m intensity in d40 (left) and d70 (right) neurons during 1 min of imaging in one FOW. Colored lines are Ca^2+^ traces of individual neurons; black line is the averaged GCamp6m signal. **l, m,** Quantification of amplitude and frequency of spontaneous individual Ca^2+^ spikes (l) and rate of synchronous firing per min of imaging in each FOW (m) (n = 10 FOW per time point from 2 independent experiment). **n,** Representative images of neurons stained with antibodies against SYNI and MAP2. **o,** Heatmap for the normalized temporal expression of transcripts important for neuronal excitability and connectivity from RNAseq experiments (n = 3 independent experiments). Data are represented as mean ± s.e.m. Dots represent individual neurons in (e, h, l) and FOW in (m). Scale bars are 50 μm (b); 100 μm (d, j); (n) 50 μm (left) and 20 μm (right). Histograms depict mean ± s.e.m. ****p < 0.0001; ***p < 0.001; **p < 0.01; *p < 0.05. Two-tailed unpaired t-test (e, h, m). Welch’s one-way ANOVA with Games-Howell’s multiple comparisons test (l).

There is a strong correlation between the date of birth and the molecular identity of cortical neurons^6, 7, 28^. Accordingly, the induction of synchronized neurogenesis generated a nearly pure cohort of early born neurons that expressed the lower layer marker TBR1 (Fig. 1b, Extended data Fig.1e, f). This contrasts with other cortical neuron culture systems in which neurogenesis occurs spontaneously over extended time periods, generating multiple neuronal identities that coexist with neural precursors in the same culture (e.g., TBR1 and SATB2 neurons in brain organoids, Extended data Fig.1g). Thus, induction of synchronized neurogenesis provides an ideal platform to selectively derive a homogeneous population of roughly coetaneous human cortical neurons and to trace their intrinsic phenotypic maturation over several months of *in vitro* differentiation.

### hPSC-derived neurons follow a gradual functional and molecular maturation program

We first tracked the morphometric development by infecting NPC at d20 with low-titer lentiviral vector encoding dTomato reporter and digitally reconstructed the morphology of individual neurons at d25, 50, 75 and 100 of differentiation (Fig.1d). We observed a significant increase in the total length of neurites over time and in the complexity of their arborization measured by Sholl analysis (Fig. 1d-f). The growth of neuronal morphologies was accompanied by the progressive acquisition of intrinsic electrophysiological properties as measured by whole-cell patch-clamp recording. Newly born immature neurons at d25 exhibit abortive or low-amplitude evoked single action potentials (AP) and over time progressively acquired more mature intrinsic functional features, including membrane potential hyperpolarization and decreased input resistance; elicited repetitive evoked AP with increased amplitude and faster kinetics (Fig.1g, h; Table S1) and developed functional excitatory synapses, as evidenced by the presence of miniature excitatory postsynaptic currents (Fig.1i). We further assessed the functional maturation at large scale through imaging of spontaneous Ca^2+^ transients in developing neurons expressing the optical calcium sensor GCaMP6m (Fig.1j, k). Consistent with the patch-clamp recordings, we observed a marked but gradual temporal increase in the amplitude and frequency of spontaneous Ca^2+^ spikes at single-neuron level (Fig.1l). Furthermore, these experiments showed a switch in network activity from sparse-to-synchronous repetitive firing by d60 of differentiation (Fig.1k, m; Movie S1,2), indicating that isochronic hPSC-derived cortical neurons, maintained in monoculture, establish synaptic connectivity and engage in synchronous network activity.

The gradual onset of functional properties correlated with the establishment of synaptic contacts as suggested by the progressive subcellular localization of SYN1 in putative presynaptic puncta-like structures (Fig.1n). This was further supported by the concerted increase in the expression of a core set of genes involved in neuronal functionality that we tracked by RNAseq at d25, 50, 75 and 100 stages (Fig.1o). These include voltage-gated Na^+^, K^+^ and Ca^2+^ channels, Na^+^/K^+^ and Ca^2+^ ATPase which regulate neuronal excitability through transmembrane electrochemical gradients and the cation/chloride transporters *KCC2* and *NKCC1* which regulate intracellular chloride homeostasis and the excitatory-to-inhibitory GABA switch during development^29^. Transcripts involved in pre- and post-synaptic compartments assembly, including SNARE, Neuroligin, Neurexin and Shank gene families and receptors for the main neurotransmitters were upregulated as well, including a *GRIN2B/A* switch in glutamate receptor subunits, known to correlate with neuronal maturity^30^.

Principal Component Analysis (PCA) of RNAseq data showed a sample distribution according to developmental stage, with hPSC-to-NPC and NPC-to-neurons representing the most molecularly distant developmental transitions. At neuron stage, more pronounced changes occurred between d25 and d50, followed by more subtle differences between d50, 75 and 100 neurons (Fig.2a). Based on PCA we inferred d25, 50 and 100 as 3 discrete maturation stages and focused on changes occurring among them. Gene Set Enrichment Analysis (GSEA) in d50 *vs.* d25 pairwise comparisons revealed a complex signature. Gene ontologies (GO) related to neuronal excitability and synaptic assembly were among the most significant. Metabolic processes, including oxidative phosphorylation, glycerolipid metabolism and PPAR signaling pathways were also significantly enriched, which points to a link between metabolism and neuronal maturation^31^. We additionally observed immunity-related GO such as antigen processing and presentation as part of the maturation program (Fig.2b). Similar GO categories were enriched in d100 *vs.* d50 comparisons (Extended data Fig.3a), further supporting the notion of a lengthy gradual unfolding of the same transcriptional signature over time. We then mapped dynamic trends of gene-expression and unbiasedly selected differentially expressed transcripts that showed monotonically increased expression during maturation (Fig. 2c and Material and Methods). Monotonically upregulated transcripts captured the molecular machinery underlying the phenotypic complexity of the maturation process, including components of the cytoskeleton (*TUBA4A, NEFH*), Ca^2+^ signaling/homeostasis (*ATP2B4*), ATP biosynthesis (*ALDOC*), lipid and cholesterol metabolism (*APOL2, NCEH1*), protein biosynthesis/degradation (*AARS, FBXO2*, *USP45*), antioxidant responses (*OXR1*), immunological changes (*HLA-B/C*) and activity-dependent transcripts (*FOS, LINC00473*) among others (Fig.2c). We validated the stage-specific expression for a selected set of monotonically upregulated markers by immunofluorescence, including *HLA-ABC*, *NEFH* and *c-FOS* (Fig.2d). The progressive upregulation of specific transcripts in our data generally matched trends of *in vivo* gene-expression based on BrainSpan Atlas of the Developing Human Brain^32^, with more pronounced changes occurring at late perinatal/early postnatal stages of human cortical development (Extended data Fig.3b).

**Fig.2:**
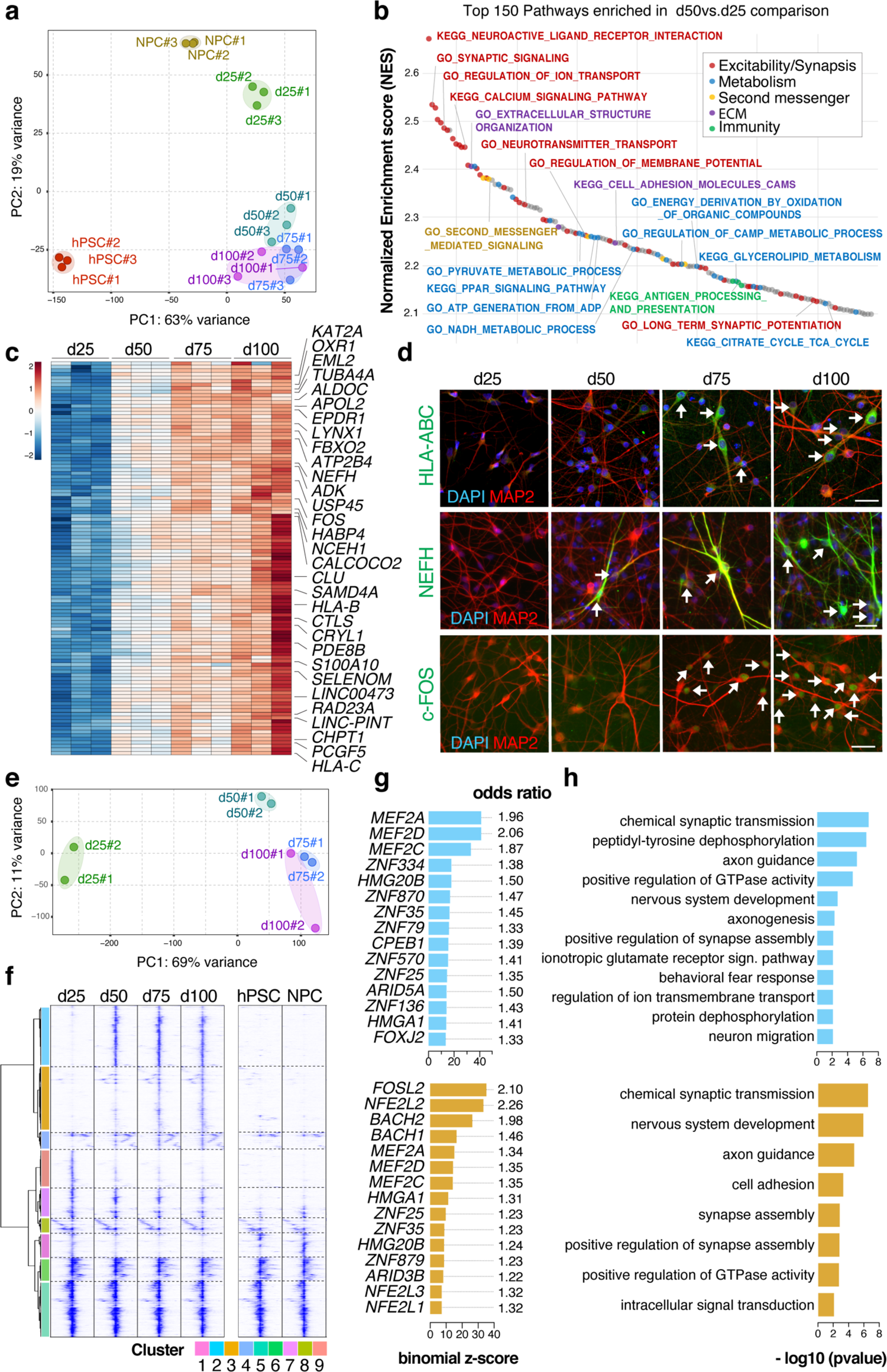
Molecular staging of neuronal maturation. **a**, PCA plot of RNAseq dataset show distribution of samples according to their time of differentiation based on top 1000 differentially expressed transcript with Variance Stabilized Normalization (VST). **b,** Waterfall plot of top 150 enriched pathways in GSEA that are enriched in more mature neurons in d50 *vs.* d25 comparison. Color codes indicate neuronal excitability/synaptic connectivity, metabolism, second messenger signaling, extracellular matrix (ECM) and immunity-related pathways. **c,** Heatmap for the normalized temporal expression of strict monotonically upregulated transcripts (maximum logFC>1, maximum RPKM>5 and s.e.m. at d100<1). **d,** Representative images of neurons at indicated time-points stained with antibodies for indicated maturation markers. **e,** PCA plot of ATACseq dataset show distribution of samples according to their maturation stage. **f,** Agglomerative hierarchical clustering by Ward linkage of differentially accessible ATACseq peaks in neurons identify 9 groups of peaks with stage-specific accessibility. **g,** Top 15 enriched transcription factor motifs at late-opening ATACseq peaks by the hypergeometric test (top, group 2; bottom, group 3). Odds ratio indicates the normalized enrichment of TF motifs in the cluster compared to the background. **h,** GO for genes linked at late-opening group 2 (top) and 3 (bottom) peaks show enrichment for synaptic-related pathways. RNAseq and downstream analyses, n = 3 independent experiments. ATACseq and downstream analyses, n = 2 independent experiments.

We next addressed the global chromatin accessibility landscape of maturation states through ATACseq at d25, 50, 75 and 100 stages. Consistent with the transcriptional data, ATACseq samples aligned along the maturation timeline by PCA (Fig.2e) with many peaks changing accessibility between d25 and d50 followed by robust but more subtle differences afterwards (Extended data Fig.4a). To uncover temporal dynamics of chromatin rearrangement, we compiled an atlas of ∼ 20,000 ATACseq peaks that included all differentially accessible peaks during neuron maturation based on pairwise comparisons of d25, 50, 75 and 100 samples. Unbiased clustering of these ATACseq peaks outlined 9 groups (Fig.2f). Except for group 5, 6 and 8, whose peaks mapped primarily at gene-promoters and showed more subtle trends of accessibility, all the other groups mapped primarily at putative enhancer DNA sequences and showed marked stage-specific dynamics of chromatin accessibility (Fig.2f; Extended data Fig.4b). Two basic patterns unfolding in a stepwise manner emerged to underlie the rewiring of the chromatin during maturation. Groups 1 and 9 defined a subset of peaks with increased accessibility in immature neurons that become less accessible towards more mature stages and that were in part shared with NPCs and hPSCs stages (Fig.2f). Conversely, peaks in groups 2, 3 and 4 showed progressive gain in accessibility and captured chromatin states of neuronal maturity. We next applied transcription factor (TF) motif enrichment analysis to infer upstream regulatory elements at the group-specific peaks. Peaks with high accessibility in young neurons were enriched for TF motifs important at early stages of cortical development^33^, including *OTX2, SOX4, EMX2, LHX2, POU3F1* and *POU3F2* among others (Extended data Fig.4c). Group 2 and 3 peaks were instead highly enriched for TF binding motifs for the Myocyte Enhancer Factor gene family (*MEF2A, D, C*) which regulate synaptic connectivity in an activity-depended manner^34^ and basic leucin zipper (bZIP) proteins such as *NFE2L2* and member of the *AP-1* complex *FOSL2*, which participate in maintaining oxidative homeostasis and proteostasis^35^ and in activity-dependent gene expression and chromatin remodeling respectively^23, 36^ (Fig.2g). We confirmed enrichment for *MEF2* and *AP-1* TF motifs (including *FOS, JUNB* and *JUND*) at late-opening differentially accessible peaks in d50 *vs.* d25 and d100 *vs.* d50 comparisons (Extended data Fig.4d). It is worth noting that the opening of peaks associated with activity-dependent TF is paralleled by their increased expression (Fig.2c, d; data not shown) and coincides with the onset of synchronous firing of the neuronal network (Fig.1j-m). Furthermore, in line with the transcriptional dynamics, genes linked to late-opening group 2 and 3 ATACseq peaks were strongly enriched for synaptic-related GO (Fig.2h). Altogether, these results indicate that synchronized hPSC-derived cortical neurons undergo a lengthy functional and molecular program of maturation and suggest their engagement in activity-dependent mechanisms at late maturation stages.

### An epigenetic switch drives neuronal maturation

We next sought to identify mechanisms responsible for the protracted pace of hPSC-derived neuron maturation. Interestingly, when focusing on genes downregulated during maturation, chromatin organization and epigenetic-related pathways emerged as the most significantly regulated by GSEA, for both d50 *vs*. d25 and d100 *vs.* d50 comparisons (Fig.3a, Extended data Fig.5a). We then specifically focused on epigenetic factors and identified a core set of transcripts whose levels monotonically decreased during maturation (Fig.3b), following a trend that largely recapitulated the expression dynamics of the same set of genes in the human cortex *in vivo* (Extended data Fig.5b). Monotonically downregulated chromatin regulators comprise members of multiple epigenetic complexes including Polycomb repressive complex 1 and 2 (PRC1/2), mammalian SWI/SNF family chromatin remodelers (BAF), MOZ/MORF acetyltransferases, nucleosome remodeling and deacetylase (NuRD), and histone lysine demethylases and methyltransferases. These results pointed to an inverse correlation between expression of a distinct set of epigenetic changes and the neuronal maturation state, raising the hypothesis that the retention of an epigenetic signature in neurons may limit their progression towards cell maturity. To this end, we performed CRISPR/Cas9 loss-of-function studies for 21 selected genes that comprise 18 chromatin regulators and the 3 transcription factors *SOX4*, *SOX11* and *KLF12* that also showed monotonic decreased temporal expression (Fig.3c). We knocked-in a Cas9 allele into the housekeeping *GPI* locus (*GPI::Cas9* hPSC line) which drives sustained Cas9 expression throughout neuronal maturation without silencing, and induced gene knock-out upon lentiviral transduction of gRNAs at neuron stage (Extended data Fig.6a-e). We differentiated *GPI::Cas9* hPSC and transduced synchronized postmitotic cortical neurons at d25 with an arrayed library of lentiviral vectors encoding dTomato reporter and gene-specific gRNAs (2 gRNAs/gene and 2 non-targeting control gRNAs; Table S2). We screened each perturbation for the ability to trigger precocious expression of cytoskeleton and pre-synaptic proteins *NEFH* and *STX1A* respectively as surrogate markers of neuronal maturity (Fig.2c, d; Extended data Fig.7b). Western Blot analysis (WB) at d35 showed increased expression of those markers across many of the gene perturbations compared to non-targeting control gRNAs (Fig.3d; Extended data Fig.7c). Furthermore, loss-of-function of about half of the perturbed chromatin regulators induced significant enhancement of neuronal functionality such as increased amplitude of individual spontaneous Ca^2+^ spikes (Fig.3e) with a trend towards increased synchronous firing rates compared to control conditions (Extended data Fig.7d). Those chromatin regulators comprise multiple PRC1/2 -related genes (*CBX2, RNF2, EPC1, EPC2, EZH2*, *MTF2*), the NuRD complex member *CHD3*, the lysine methyltransferase *KMT5B,* the BAFs members *SMARCA4* and *SMARCAD1* and the bromodomain containing gene *BRD1*. While our study does not rule out that other gene categories identified in our study (Fig.2b) or additional epigenetic factors not validated here (Fig.3b), may further regulate the maturation process, we have identified a set of chromatin regulators whose loss-of-function can robustly accelerate the onset of molecular and functional maturation features. Importantly, this epigenetic signature is similarly downregulated during the maturation of multiple cortical neuron subtypes in the developing mouse cortex *in vivo* (Fig.3f, g) albeit at a much more rapid pace than in human cells. Thus, the progressive downregulation of key chromatin regulators during neuronal maturation combined with their loss-of-function data suggests the existence of an epigenetic “brake” that is released gradually to ensure the protracted unfolding of maturation programs in human neurons.

**Fig.3:**
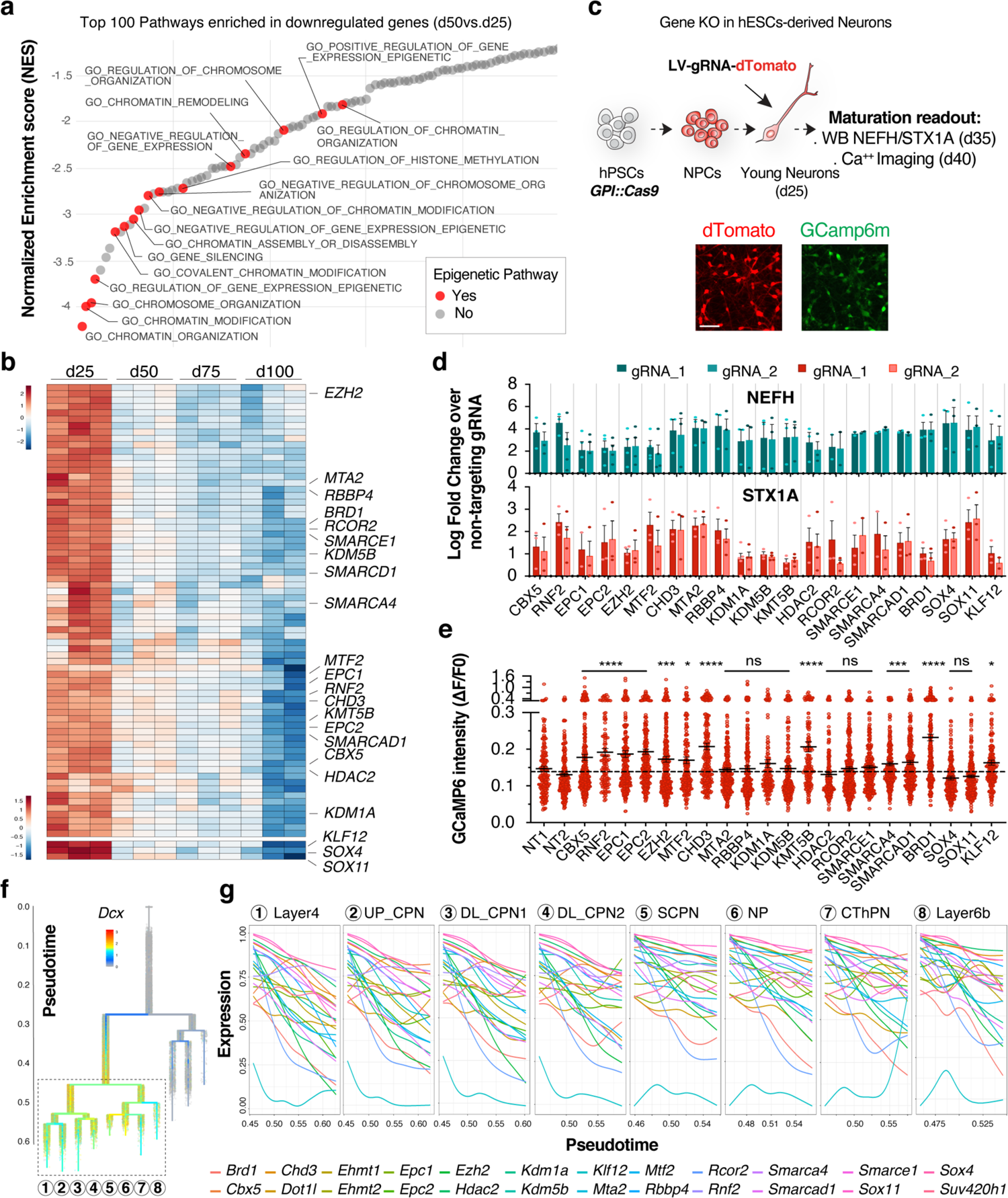
An epigenetic switch drives neuronal maturation. **a**, Waterfall plot of GSEA enriched pathways from RNAseq studies that are negatively correlated with neuronal maturation in d50 *vs.* d25 comparison. Red dots indicate epigenetic-related pathways. **b,** Heatmap for normalized temporal expression of chromatin regulators that are monotonically downregulated during maturation (maximum logFC > 1, s.e.m. at d100 <1). Gene labeled in the heatmap were selected for perturbation studies. **c,** Schematic of experimental paradigm for gene-KO in postmitotic hPSCs-derived neurons: Cas9 expressing neurons at d25 were infected with lentiviral vectors encoding gene-specific gRNAs. Induction of precocious molecular and functional maturation was assessed by WB and Ca^2+^ imaging respectively. **d,** WB analysis for the expression of NEFH and STX1A maturation markers upon gene-KO in neurons (2 gRNA/gene). Histograms depict average log_2_Fold change ± s.e.m. over neurons transduced with non-targeting gRNA. Dots represent replicates (n = 3 independent experiments). **e,** Amplitude of spontaneous Ca^2+^ spikes of individual neurons in gene-KO experiments. Dotted line represents the average amplitude of the 2 non-targeting gRNA controls and dots represent individual neurons (n = 3-6 FOW from 2 independent experiments). (a, b) n = 3 independent experiments. **f,** Branching tree from single-cell RNAseq from Di Bella et al.^33^ showing expression of *Dcx* transcripts in the mouse neocortex. **g**, Temporal expression of chromatin regulators from perturbation studies in hPSC-derived neurons (d) in multiple neuronal identities in the mouse neocortex. UP, upper layer; DL, lower layer; CPN, callosal projection neurons; SCPN, subcerebral projection neurons; NP, near projecting; CThPN, cortico-thalamic projection neurons. ****p < 0.0001; ***p < 0.001; **p < 0.01; *p < 0.05. Welch’s one-way ANOVA with Games-Howell’s multiple comparisons test.

### An epigenetic barrier in NPCs sets the rate of human neuronal maturation

Temporal expression analysis revealed that most of the epigenetic factors identified as regulators of neuronal maturation in our loss-of-function studies (hits) were already expressed already in dividing NPC (Fig.4a). This raised the intriguing possibility that those chromatin regulators may establish an “epigenetic barrier” at the hPSC-to-NPC transition, well before the onset of neurogenesis. To test this hypothesis, we manipulated a subset of the epigenetic hits directly at the NPC stage, and we selected small molecule inhibitors of the PRC2 regulator *EZH2*, the histone lysine methyltransferases *EHMT1/2*, *DOT1L*, *KMT5B* and demethylases *KDM1A* and *KDM5* (Extended data Fig.8a, b). We treated NPCs only transiently, following the induction of cortical CNS patterning from d12 to d20. The small molecules were washed out and withdrawn starting at d20, prior to the induction of synchronized neurogenesis. Neurons derived from NPC treated with the epigenetic inhibitors versus DMSO were grown under the same exact culture conditions (Fig.4b). Importantly, neither treatment condition altered the expression of *PAX6* or *FOXG1* and there was no evidence of precocious neurogenesis based on the ratio of Ki67^+^ NPC and Map2^+^ neurons at d20 (Extended data Fig.8c, d). We next assessed the extent of maturation induced by WB and Ca^2+^ imaging at d35 and d40 respectively (Fig.4b). Among the treatments, transient inhibition of *EZH2*, *EHMT1/2* and *DOT1L* in NPC with GSK343, UNC00638 and EPZ004777, respectively, induced robust increased expression of the maturation markers NEFH and STX1A in d35 neurons (Fig.4c; Extended data Fig.9a). In addition, transient *EZH2* inhibition in NPC triggered a marked and highly significant increase in all measured functional properties such as amplitude and frequency of individual Ca^2+^ spikes and synchronicity of the neuronal network respect to DMSO control neurons (Fig.4d-f; Movie S3,4). *EHMT1/2* inhibition induced significant changes in the amplitude and synchronicity of Ca^2+^ spikes while neurons derived from NPC treated with the *DOT1L* inhibitor showed a modest functional enhancement and a trend towards increased synchronous firing rates (Fig.4d-f; Movie S3,5,6). We next assessed global transcriptional changes in neurons derived from NPC treated with either EZH2, EHMT1/2 or DOT1L inhibitors by RNAseq at d35, which revealed robust changes in all treated conditions respect to DMSO controls (Extended data Fig.9b-c). Downregulated genes captured pathways typically found in NPC and newborn neurons (Fig.4g), including members of the Sox family of TF and Notch pathway-related transcripts among others (data not shown). In line with the enhanced functionality of neurons derived from treated NPC, upregulated transcripts were robustly enriched for maturation related GO, including chemical synaptic transmission and ion transmembrane transport (Fig.4g). These results demonstrated that enhancement of maturation state can be achieved through transient inhibition of chromatin regulators at the NPC stage and identified EZH2, EHMT1/2 and DOT1L as key components of the epigenetic barrier.

**Fig. 4:**
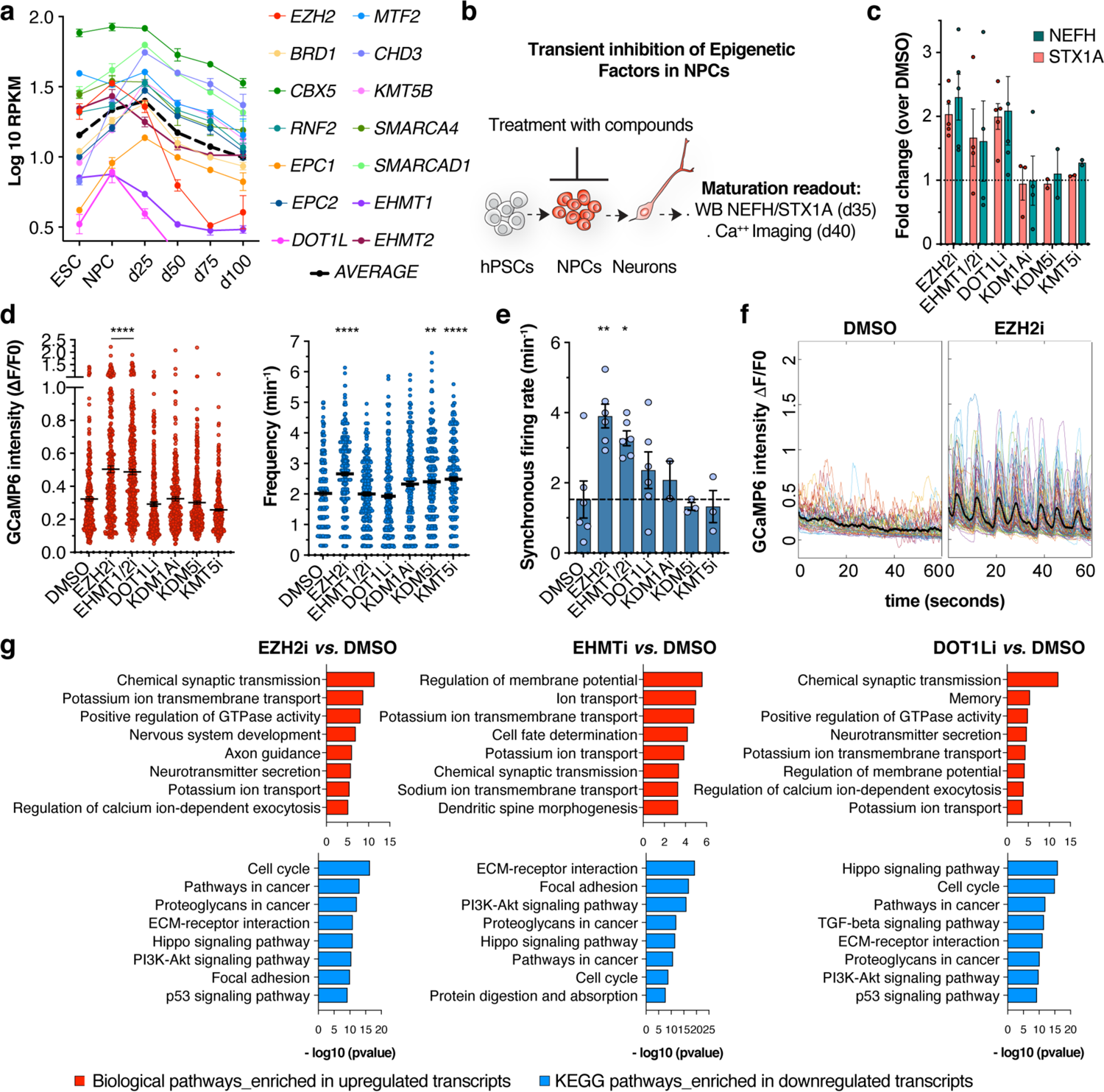
Transient inhibition of epigenetic factors in NPCs drive faster maturation in neurons. **a**, Temporal expression of chromatin regulators hits from gene-KO studies at hPSCs, NPCs and neuron stages (n = 3 independent experiments). **b,** Schematic of experimental paradigm for transient inhibition of chromatin regulators at progenitor cell stage. NPCs were treated with small molecule from d12 to d20. Control and treated NPCs were induced for synchronized neurogenesis and neurons derived from all the treatments were maintained in the same conditions. Induction of preconscious molecular and functional maturation was assessed by WB and Ca^2+^ imaging respectively. **c,** WB analysis for the expression of NEFH and STX1A maturation markers. Histograms depict average fold change ± s.e.m. over DMSO controls. Dotted line represents DMSO controls. Dots represent replicates (n = 2-5 independent experiments). **d-e,** Amplitude and frequency of spontaneous Ca^2+^ spikes of individual neurons (d) and synchronicity rate of spontaneous network activity (e) in treated *vs*. control conditions. Dots represent individual neurons in (d) and FOW in (e) (n = 3-6 FOW from 2 independent experiments). **f**, Representative traces of normalized GCamp6m intensity in DMSO control (left) and EZH2i (right) conditions during 1 min of imaging in one FOW. Colored lines are Ca^2+^ traces of individual neurons while black line is the averaged GCamp6m signal. **g**, GO for upregulated (top) and downregulated (bottom) transcripts from RNAseq studies in pairwise comparisons (n = 3 independent experiments). ****p < 0.0001; ***p < 0.001; **p < 0.01; *p < 0.05. Welch’s one-way ANOVA with Games-Howell’s multiple comparisons test (e); ordinary one-way ANOVA (f).

To gain insights into the epigenetic regulation of maturation programs, we characterized the dynamics of H3K27ac, H3K4me3, H3K27me3 and H3K9me3 histone post-translational modifications (PTMs) in hPSC-derived cortical NPC and neurons via CUT&RUN experiments. Unsupervised clustering of CUT&RUN peaks with differential binding for histone PTMs in NPC *vs.* Neurons identified 8 groups of peaks characterized by distinct combinatorial patterns of histone PTMs (Extended data Fig.10a). GO analysis on genes linked to CUT&RUN peaks identified cluster 1, 2 and 3 as highly enriched for chemical synaptic transmission and ion transmembrane transport terms among others, indicating that epigenetic regulation at these genomic loci may play a role in driving maturation-related gene expression (Extended data Fig.10c). We then intersected the genes linked to each CUT&RUN cluster with all differentially expressed genes by RNAseq in NPC, d25, d50, d75 and d100 neurons, irrespective of the directionality of the changes. This analysis identified a correlation between the patterns of histone PTMs in clusters 1, 2 and 3 and maturation-dependent changes in gene expression (Extended data Fig.10e). Furthermore, maturation-related genes belonging to cluster 1, 2 and 3 were among the statistically upregulated transcripts in neurons derived from NPC transiently treated with DOT1L and EZH2 inhibitors versus those derived from DMSO treated control NPC (Extended data Fig.10f). Cluster 2 defined a subset of peaks with increased dual binding for H3K27ac and H3K4me3 histone PTMs at neuron stage, marking active chromatin domains at putative enhancer sequences, enriched for activity-dependent TF motif such as *AP1* and *MEF* gene families (Extended data Fig.10b, d) suggesting that activity-dependent mechanisms contribute to driving neuronal maturation. In contrast, cluster 1 is dominated by the dual presence of the EZH2 dependent H3K27me3 repressive mark and the active H3K4me3 PTM at NPC stage. Such poised or bivalent state^37^ is resolved toward active chromatin state at neuron stage via loss of the repressive H3K27me3 mark and acquisition of modest levels of the active H3K27ac PTM (Extended data Fig.10a; Fig. 5a). Cluster 3 showed a similar pattern with a partial bivalent state in NPC and a more pronounced acetylation of H3K27 in neurons (Extended data Fig.10a). These results indicate a key role for the EZH2 dependent deposition of the H3K27me3 repressive mark in maintaining maturation programs in a poised state, a finding further supported by the increased expression of various maturation genes (with bivalent chromatin state in NPC; Fig.5a, b) upon transient treatment with EZH2 inhibitors. Those transcripts match the unperturbed, chronological maturation signature (Fig.1o; Fig.2b-d) and are involved in synaptic assembly and functionality, activity-dependent mechanisms (*FOS, FOSB, NPAS4, BDNF*), glycerolipid metabolism and PPAR signaling (*DGKK, DGKG, PPARG)*, maturation of the cytoskeleton (*NEFH, TUBA4A*) and immunological programs (*HLA-B* and *C*) (Fig.5b, c). Interestingly, among the bivalent genes at NPC stage, we found several chromatin regulators that show a gradual increase in expression during cortical neuron maturation that were induced dose-dependently in NPC upon transient epigenetic inhibition (Fig.5d-f). These include *JADE2* (also known as *PHF15*), a ubiquitin ligase that target for degradation *KDM1A*^38^, whose loss-of-function triggered increased expression of maturation markers (Fig. 3d) and *CHD5*, which facilitates the expression of neuron specific gene programs^39^. These results indicate that the epigenetic barrier identified in our study controls the temporal onset of maturation via a dual mechanism; by directly maintaining maturation genes in a poised state and indirectly by modulating the expression of competing epigenetic regulators promoting maturation (Fig 5g, h).

**Fig.5:**
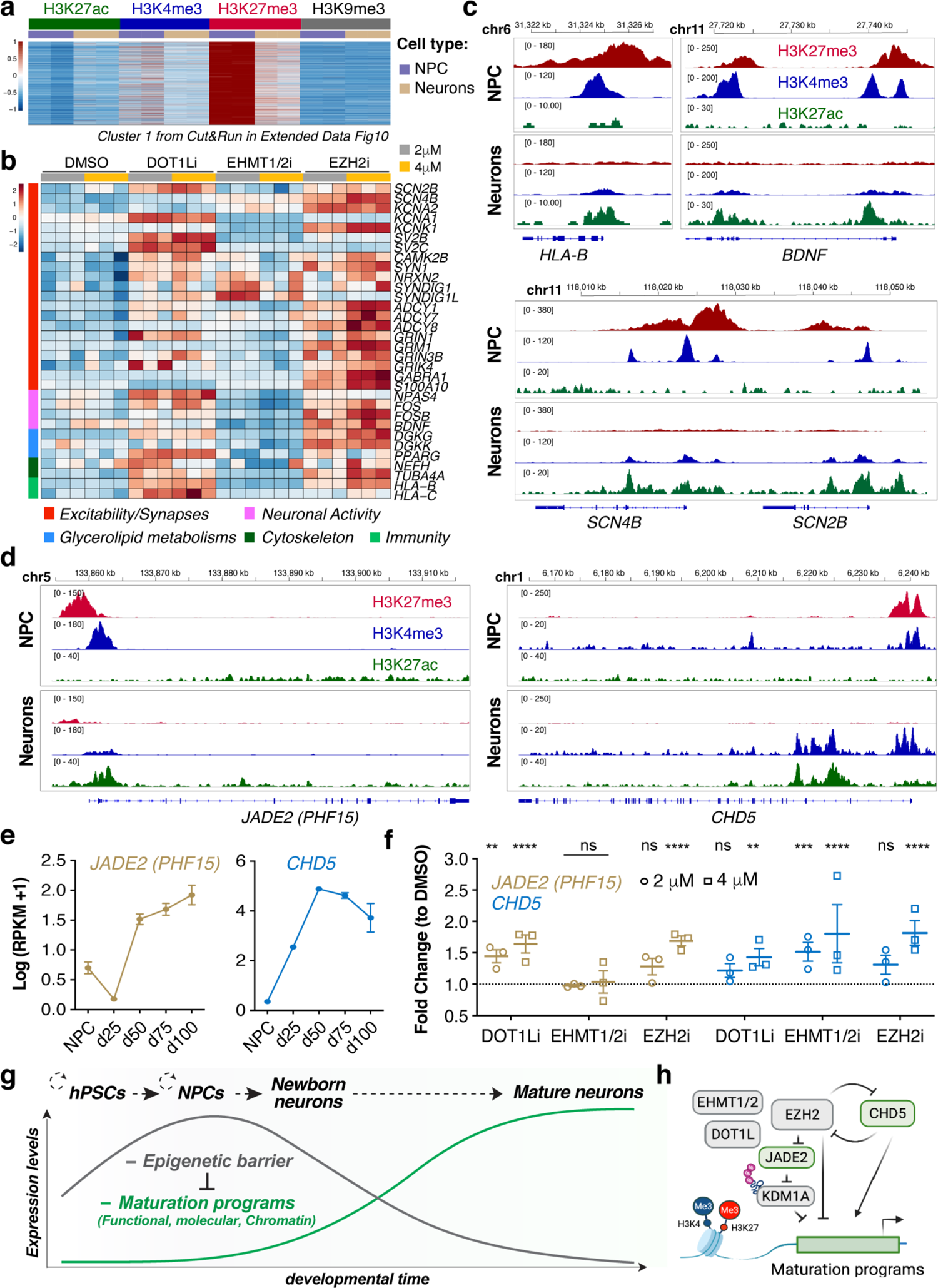
An epigenetic barrier in NPCs control the onset of maturation programs. **a**, Heatmap for cluster 1 from Extended data Fig.10 representing bivalent peaks in NPCs decorated by H3K27me3 and H3K4me3 that lose H3K27me3 at neuron stage (n = 2 replicate for each condition). **b,** Heatmap for the normalized expression of representative transcripts within the bivalent genes in d35 neurons derived from NPC treated with DOT1l, EHMT1/2 and EZH2 inhibitors at 2 and 4 μM (n = 3 replicate for each condition). **c-d,** Representative Integrative Genome Viewer density tracks for H3K27me, H3K4me3, and H3K27ac at indicated genomic loci in NPC and neurons (n = 2 replicate for each condition). **e,** Temporal expression of *CHD5* and *JADE2* transcripts through the maturation time-course (n = 3 independent experiments). **f,** Expression of *CHD5* and *JADE2* transcripts in d35 neurons derived from NPC treated with DOT1l, EHMT1/2 and EZH2 inhibitors at 2 and 4 μM (n=3 replicate for each condition). Histograms depict mean ± s.e.m. **g-h,** Schematic of the main conclusion of the study. **g**, The temporal unfolding of maturation signatures in hPSC-derived neurons proceed gradually and is marked by the retention of multiple epigenetic pathways that establish an epigenetic barrier at progenitor cell stage that gets inherited in neurons. **h,** Key members of the epigenetic barrier, including EZH2, maintain maturation programs in a poised state through deposition of repressive histone marks.

## Discussion

Novel paradigms in stem cell biology provide new approaches to investigate mechanisms regulating the timing of developmental transitions. Here, we have developed a method that synchronizes the generation of cortical neurons from hPSC and facilitates the study of neuronal maturation at high precision. Using this platform, we established a framework for the maturation of hPSC-derived cortical excitatory neurons and dissected an epigenetic mechanism responsible for their protracted maturation. While we focused on early born excitatory cortical neurons, our strategy could be readily applied to other relevant hPSC-derived lineages to investigate shared and unique signatures of maturation across cell types and the impact of cell-to-cell interaction on the maturation process. In our study, we specifically addressed neuron intrinsic mechanisms and developed a culture system that is largely devoid of glial cells such as astrocytes, which are known to impact neuronal functionality^21, 22^. It would be interesting to address complementary aspects of the maturation program that are modulated by neuron/glia interactions and by others extrinsic environmental factors. Indeed, our study pointed out multiple phenotypic dimensions underlying the neuronal maturation process, including synaptic, metabolic, immunity-related, and epigenetic programs that unfold gradually in hPSC-derived neurons. It is presently unclear whether the temporal onset for each of those aspects is controlled by multiple, phenotype-specific mechanisms or whether a shared clock-like mechanism orchestrates their concerted expression. An example of a specific, synapse-related mechanism is the emergence of human paralogs for the postsynaptic *SRGAP2A* gene, which have been shown to negatively regulate the ancestral gene and that is associated with human-specific protracted synaptogenesis and dendritic spines morphogenesis^40, 41^. Similarly, human-specific duplication of the Notch family gene *NOTCH2NL* has been linked to an extended expansion of progenitor cells characteristic of the human cortex via increase of the NOTCH signaling pathway^42, 43^. These results suggest that the evolutionary divergence of developmental timelines may have emerged in part through modulation of ancestral processes.

In our study we identified a distinct epigenetic signature (i.e., epigenetic barrier) that is gradually downregulated in developing neurons and that prevents precocious onset of the multiple dimensions of maturation programs. Our data indicate that expression levels and rates of downregulation of components of the epigenetic barrier determine the pace at which maturation unfolds, ensuring a protracted timeline in human neurons. The epigenetic barrier identified here comprises multiple families of chromatin regulators, and further studies will be needed to precisely dissect the interplay among the factors identified. For example, we observed that *SOX4* and *SOX11* TF showed a gradual downregulation similar to the epigenetic barrier and could trigger precocious expression of maturation markers upon loss-of-function. Interestingly, *Sox4* and *Sox11* follow similar temporal trends in the mouse cortex^44^ and distinct dynamics in *Sox11* downregulation have been recently shown to impact the pace of maturation of mouse cortical neurons *in vivo*^45^, further reinforcing the importance of gene expression dynamics in controlling neuronal maturation rates. Recent studies have correlated differences in the rate of fundamental biological processes, including transcription initiation and protein turnover, with species-specific timescale of early development^11, 12, 14^. However, further studies are needed to infer causal relationships and to elucidate whether those mechanisms similarly apply to the later and much more protracted process of neuronal maturation. It will be interesting to determine whether the epigenetic switch identified in our study represents a complementary or parallel mechanism driving human-specific timing.

A central finding of our study is the discovery that the epigenetic barrier at maturation is established before the onset of neurogenesis during hPSC-to-NPC transition and in turn gets inherited in newborn neurons. Our data indicate that the rate of neuronal maturation is already “pre-programmed” at the precursor cell stage as the transient inhibition of key epigenetic factors such as EZH2, DOT1L and EHMT1/2 prior to neurogenesis is sufficient to prime postmitotic neurons for rapid acquisition of functionality. Furthermore, focusing on EZH2, the catalytic subunit of PRC2, we found that several maturation-related genes are transcriptionally poised at the NPC stage through deposition of the H3K27me3 repressive histone mark at their promoter. Poised genes under EZH2-mediated silencing via H3K27me3 comprised “effectors” of neuronal maturity (e.g., ion channels and synaptic proteins) and a subset of chromatin regulators that has been shown to functionally interact with the components of the epigenetic barriers, such as the *KDM1A* ubiquitin-ligase *PHF15* (*JADE2*)^38^ and *CHD5,* which facilitates neuronal gene expression^39^ and mutually crosstalk with EZH2^46, 47^. Thus, our results suggest a complex interplay among different epigenetic factors and future studies will have to further elucidate the molecular interactions and hierarchy of this epigenetic regulatory network modulating neuronal maturation. Interestingly, EZH2 has been recently proposed to epigenetically drive the temporal progression and neurogenic competence of NPC in the developing mouse cortex^48, 49^. Similarly to our study on cortical neuron maturation, the downregulation of EZH2 in cortical NPC correlated with the increased expression of maturation-related GO in precursor cells^48^, raising the interesting possibility that the same epigenetic machinery regulates the progression of NPC and the maturation of post-mitotic neurons in order to synchronize the timelines of brain development.

## Acknowledgments

We thank members of the Studer lab for discussion and insights into the project. We also thank Tao Tong and Christina Pirgaky from the Bio-Imaging Resource Center at Rockefeller University for guidance in the design and analysis of Ca^2+^ imaging experiments. We would like to thank the Epigenomics Core at Weill Cornell Medical College, the Epigenetic Innovation lab, and the Integrated Genomic Operation (IGO) core at MSKCC.

## Author contributions

G.C.: conceptualization, study design, performing most of the wet lab experiments, data analysis and interpretation, writing of manuscript. L.S.: conceptualization, study design, data interpretation, writing of manuscript. H.C., M.K. and R.P.K: computational analyses. C.S.L. and R.P.K: supervision of computational analysis and interpretation of data. A.B.: Western Blots, CUT&RUN, data analysis and interpretation. K.A.A: performed electrophysiological recordings and interpretation of data, P.A.G.: supervision of electrophysiological recordings and interpretation of data. R.M.W. organoid culture, design of GPI locus targeting, computational analysis. All authors provided feedback in editing the manuscript.

## Material & Correspondence

WA-09 lines created (GPI::Cas9) will be made available upon request from the Studer lab at MSKCC under a Material Transfer Agreement with the institute.

## Data Availability

All genomic datasets have been deposited at GEO (accession number pending). Publicly available datasets of human brain development were from BrainSpan atlas of the developing human brain (https://www.brainspan.org). Published dataset of mouse cortical development were from Di Bella et al., Nature 2021; PMID: 34163074^33^.

## Methods

### hPSCs cell lines and culture

Work using human pluripotent stem cells (hPSC) was approved in compliance with the Tri-Institutional ESCRO at Memorial Sloan Kettering Cancer Center, Rockefeller University and Weill Cornell Medicine. hPSC WA09 (H9; 46XX) and derivate GPI::Cas9 were maintained with Essential 8 media (Life Technologies #A1517001) in feeder-free conditions onto Vitronectin (VTN-N, Thermo Fisher #A14700) coated dishes. hPSCs were passaged as clumps every 4-5 days with EDTA (0.5M EDTA/PBS) and routinely tested for mycoplasma contamination. GPI::Cas9 knock-in hPSCs line was generated using CRISPR/Cas9-mediated homologous recombination by transfecting H9 hPSCs with the Cas9-T2A-Puro targeting cassette downstream of the GPI gene (Fig. S6B). Selected clones were validated by genomic PCR and Cas9 mRNA and protein expression by qRT-PCR and Western Blot respectively and screened for Karyotype banding.

### Synchronized generation of cortical neurons

hPSCs (passage 40-50) were differentiated toward cortical excitatory neurons using an optimized protocol based on dual-SMAD inhibition and WNT inhibition as following. hPSCs were dissociated at single cells using Accutase and plated at 300,000 cells/cm^2^ onto Matrigel (#354234, Corning) coated wells in Essential 8 media supplemented with 10 μM Y-27632. On day 0-2, cells were fed daily by complete medium exchange with Essential 6 medium (E6, #A1516401, Thermo Fisher Scientific) in the presence of 100 nM LDN193189 (#72142, Stem Cell Technologies), 10 μM SB431542 (#1614, Tocris) and 2 μM XAV939 (#3748, Tocris) to induce anterior neuroectodermal patterning. On day 3-9 cells were fed daily with Essential 6 medium (E6, #A1516401, Thermo Fisher Scientific) in the presence of 100 nM LDN193189 (#72142, Stem Cell Technologies), 10 μM SB431542. On day 10-20 cells were fed daily with N2/B27 media (1:1 NB:DMEM/F12 basal media supplemented with 1X N2 and B27 minus vitamin A to generate a neurogenic population of cortical neuronal progenitor cells (NPCs). N2 and B27 supplements were from Thermo. At day 20, NPCs were either cryopreserved in STEM-CELLBANKER solution (Amsbio) or induced for synchronized neurogenesis as following: NPCs were dissociated at single cells following 45 min incubation with Accutase and seeded at 150,000 cells/ cm^2^ onto poly-L-ornithine and Laminin/ Fibronectin coated plates in NB/B27 medium (1X B27 minus vitamin A, 1% L-glutamine and 1% Pen/Strep in Neurobasal medium) in presence of 10 μM Notch pathway inhibitor DAPT for 10 days (until day30). For long term culture, neurons were maintained in NB/B27 supplemented with BDNF (#450-10, PreproTech), GDNF (#248-BD-025, R&D biosystems), cAMP (#D0627, Sigma) and AA (#4034-100, Sigma). From day 20 onwards, cells were fed every 4/5 days.

### EdU labelling and small molecule treatments

For birthdating experiments of hPSC-derived cortical neurons, 3μM EdU (5-ethynyl-2′-deoxyuridine, A10044 Invitrogen) was added to the culture for 48h in the following time window: day18/19, day20/21, day22/23, day24/25, day26,27, day28/29. After treatment, EdU was washed out and neurons were fixed at day40 of differentiation and processed for immunostaining. Treatment of cortical neuronal progenitor cells (NPCs) with small molecules inhibitors of chromatin regulator was performed from day12 to 20 of differentiation (Fig. 5B). List of small molecules are relative intracellular target is reported in Fig. S8B. Small molecules were dissolved in DMSO and added to the N2/B27 media at 2 or 4 μM depending on the experiment. Small molecules were washed out before the induction of synchronized neurogenesis and neurons derived from all the treatments were maintained in the same conditions.

### Morphological reconstructions

hPSCs derived neurons were infected with low titer lentiviruses expressing dTomato reporter at day 20 and fixed at day 25, 50, 75 and 100. The dTomato reporter signal was amplified by immunofluorescence staining and individual neurons were imaged at 10x. Neuronal morphology was reconstructed using the filament tracing function of Imaris software. Measurements were performed in the Imaris platform and extracted for quantifications and statistics.

### Immunofluorescence

Cultured cells were fixed with 4% PFA in PBS for 20min at RT, washed three times with PBS, permeabilized for 30 min in 0.5% Triton X-100 in PBS and then blocked in a solution containing 5% Normal goat serum, 2% BSA and 0.25% Triton X-100 for 1h at RT. Primary antibodies were incubated overnight at 4°C. The following primary antibodies were used: rabbit anti-Pax6 (901301, Biolegend); rabbit anti-FoxG1 (M227, Clonetech); mouse anti-Nestin (M015012, Neuromics); mouse anti MAP2 (M1406, Sigma); chicken anti-MAP2 (ab5392, Abcam); rabbit anti-Class III β-tubulin TUJI (MRB-435P, Covance); mouse anti-Ki67 (M7240, Dako); rabbit anti-Ki67 (RM-9106, Thermo Scientific); rabbit anti-Tbr1 (ab183032, Abcam); rat anti-Ctip2 (ab18465, Abcam); mouse anti-Satb2 (ab51502, Abcam); rabbit anti-Synapsin I (S193, Sigma); mouse anti-Neurofilament H (non phosphorylated) (SMI32; Enzo Life science); mouse anti c-Fos (ab208942, Abcam); mouse anti-HLA Class I ABC (ab70328, abcam); goat anti-RFP (200-101-379, Rockland); rabbit anti-DsRed (632496, Clontech). EdU^+^ cells were detected using the Click-iT EdU Imaging kit (Molecular Probes) with Alexa Fluor 488. Secondary antibodies conjugated to either Alexa 488, Alexa 555 or Alexa 647 (Thermo) were incubated for 45 min. Cell nuclei were stained with 5 μM 4′-6-diamidino-2-phenylindole (DAPI) in PBS1X.

### Electrophysiological recording

Neurons were plated in 35 mm dishes and whole-cell patch clamp recordings were performed at day 25, 50, 75 and 100 of differentiation as previously described ^19^. Briefly, neurons were visualized using a Zeiss microscope (Axioscope) fitted with 4x objective and 40x water-immersion objectives. Recordings were performed at 23 – 24 °C and neurons were perfused with freshly prepared ACSF extracellular solution saturated with 95% O2 – 5% CO_2_ that contained (in mM): 126 NaCl, 26 NaHCO_3_, 3.6 KCl, 1.2 NaH_2_PO_4_, 1.5 MgCl_2_, 2.5 CaCl_2_, and 10 glucose. Pipette solution for all recordings contained (in mM): 140 CsCl, 10 NaCl, 10 HEPES, 0.5 EGTA, 3 Mg-ATP, 0.2 Na-GTP, and 10 Na_2_-phosphocreatine, pH adjusted to 7.3 with CsOH. 20 μM (−)-Bicuculline methochloride (Tocris), 1 μM strychnine HCl (Sigma), and 0.5 μM tetrodotoxin (TTX) (Alomone Labs) were added to the ACSF for mEPSC recordings to block GABA_A_ receptors, glycine receptors, and Na^+^ channels, respectively. Input resistance was measured from a voltage response elicited by intracellular injection of a current pulse (−100 pA, 200 ms). Membrane voltage was low-pass filtered at 5 kHz and digitized at 10 kHz using a Multiclamp 700B amplifier connected to a DigiData 1322A interface (Axon Instruments) using Clampex 10.2 software (Molecular Devices, Foster City, CA). Liquid junction potentials were calculated and corrected off-line. Action potentials (AP) were generated in current clamp from currents injected in 10 pA intervals from 0 to 250 pA. Recordings were analyzed for: resting membrane potential, input resistance, rheobase, threshold, as well as AP amplitude, overshoot, duration, half-width, rise and decay. Neurons were held at −80 mV and continuous recordings of mEPSCs were made using Axoscope software (Molecular Devices, Union City, CA). Data processing and analysis were performed using MiniAnalysis (Synaptosoft, Decatur, GA) and Clampfit 10 (Molecular Devices). Events were detected by setting the threshold value, followed by visual confirmation of mEPSC detection.

### Calcium imaging and analysis

hPSC-derived cortical neurons were infected with lentiviruses encoding GC GCamp6m and cultured on μ-plate 96 Well Black (Ibidi). Ca^2+^ was performed as previously described ^50^. Briefly, on the day of the imaging, cells were gently washed twice in modified Tyrode solution (25 mM HEPES (Invitrogen), 140 mM NaCl, 5 mM KCl, 1 mM MgCl_2_, 10 mM glucose, 2 mM CaCl_2_, 10 μM glycine, 0.1% BSA pH 7.4, pre-warmed to 37 °C) and equilibrated in imaging buffer for 1-2 min (25 mM HEPES, 140 mM NaCl, 8 mM KCl, 1 mM MgCl_2_, 10 mM glucose, 4 mM CaCl_2_, 10 μM glycine, 0.1% BSA pH 7.4, pre-warmed to 37 °C). GCaMP6m fluorescence was recorded on Celldiscover7 (ZEISS) inverted epi-fluorescence microscope with the 488 nm filter under environmental control (37 °C; 95% O_2_ – 5% CO_2_) at the Bio-Imaging Resource Center (BIRC) at Rockefeller University. Neuronal cultures were imaged for ∼3 min at a frame rate of 4–6 frames/second (800 frames/time lapse) using a 10x or 20x objective. Analysis was performed as previously described ^50^. Briefly, the live-imaging image stack was converted to TIFF format and loaded into optimized scripts in MATLAB. Region of Interest (ROI) were placed on the neuron somas to calculate the raw GCaMP6m intensity of each neuron over time. The signal intensity of each raw trace was normalized to the baseline (ΛF/F0) for spike detection. Single-neuron amplitude was calculated from the normalized GCaMp6m intensity for all the detected spikes in each trace (mean ΔF/F0 of detected spikes for each neuron). Single-neuron frequency was calculated as the number of detected spikes in each trace per minute of recording. Network activity was assessed by calculating the synchronous firing rate, defined as the number of detected synchronous Ca^2+^ spikes from all ROI in one Field of View (FOV) per minute of recording.

### Image analysis and quantification

Morphological reconstruction of neurons was performed using Imaris Software. Ca^2+^ imaging analysis was performed using MATLAB software. Quantification of immunofluorescence images was performed in ImageJ or using the Operetta High content imaging system coupled with Harmony software (PerkinElmer).

### Protein extraction and Western Blots

Cells were harvested and lysed in RIPA buffer (Sigma) with 1:100 HaltTM Protease and Phosphatase Inhibitor Cocktail (Thermo Fisher Scientific) and then sonicated for 3×30sec at 4°C. Protein lysates were centrifugated for 15 min at > 15000 rpm at 4°C and supernatant was collected and quantified by Precision Red Advanced Protein Assay (Cytoskeleton). 5-10 μg of protein were boiled in NuPAGE LDS sample buffer (Invitrogen) at 95°C for 5 min and separated using NuPAGE 4%–12% Bis-Tris Protein Gel (Invitrogen) in NuPAGE MES SDS Running Buffer (Invitrogen). Proteins were electrophoretically transferred to nitrocellulose membranes (Thermo Fisher Scientific) with NuPAGE Transfer Buffer (Invitrogen). Blots were blocked for 60 min at RT in TBS-T + 5% nonfat milk (Cell Signaling) and incubated overnight in the same solution with the respective primary antibodies at 4°C. The following primary antibodies were used: mouse anti-Neurofilament H (non-phosphorylated) (SMI32; Enzo Life science); mouse anti-Syntaxin 1A (110 111; SYSY); mouse anti-actin (MAB1501; Millipore); mouse anti-Cas9 (1497; Cell Signaling Technology); rabbit anti-Chd3 (ab109195, Abcam); rabbit anti-KDM5B (ab181089, abcam). The following secondary antibodies were incubated for 1 hour at RT: anti-mouse IgG HRP-linked (7076; Cell Signaling Technology) and anti-rabbit IgG HRP-linked (7074; Cell Signaling Technology) Blots were revealed using SuperSignalTM West Femto Chemiluminescent Substrate (Thermo Fischer Scientific). Chemiluminescence was imaged and analyzed using Image lab software (Biorad).

### RNA isolation and qRT-PCR

Samples were collected in Trizol and total RNA was isolated by chloroform phase separation using Phase Lock Gel-Heavy tubes, precipitated with EtOH, and purified using RNeasy Mini Kit (Qiagen) with on-column DNA digestion step. cDNA was generated using the iScript Reverse Transcription Supermix (Bio-Rad) for RT-qPCR and qPCR reactions were performed using SsoFast EvaGreen^®^Supermix (Bio-Rad) using Quantitect Primer assays (QUIAGEN). Results were normalized to the housekeeping gene GAPDH.

### DNA construct and lentivirus production

A Cas9-T2A-PuroR cassette flanked by 5’ and 3’ homology arms for the GPI locus was generated by NEBuilder^®^ HiFi DNA Assembly Cloning Kit of PCR amplified fragments according to manufacturer’s instruction. EF1alpha-GCamp6m lentiviral vector was generated by PCR amplification of GCamp6m from pGP-CMV-GCaMP6m (Addgene #40754) using with Q5 High Fidelity master mix (NEB) and subcloned into pWPXLd (Addgene #12258) into BamH1 and EcoRI restriction site using standard cloning methods. For the simultaneous expression of gene-specific gRNA under transcriptional control of U6 promoter and dTomato fluorescent reporter driven by EF1alpha promoter, the SGL40.EFs.dTomato vector (Addgene #89398) was modified by inserting a P2A-Basticidin cassette downstream of dTomato sequence to generate the SGL40.EFs.dTomato-Blast backbone. gRNA sequences specific to each gene were designed using SYNTEGO CRISPR design tool (https://www.synthego.com/products/bioinformatics/crispr-design-tool) and validated using CRISPOR tool ^51^ (http://crispor.tefor.net). DNA oligos (IDT) were annealed and subcloned into BsmBI restriction sites of SGL40.EFs.dTomato-Blast lentiviral backbone by standard cloning methods. Lentiviruses were produced by transfection of HEK293T cells using the Xtreme Gene 9 DNA transfection reagent (Sigma) with the respective lentiviral vectors along with the packaging vectors psPAX2 (Addgene, 12260) and pMD2.G (Addgene, 12259). Arrayed CRISPR gRNA lentiviral libraries were produced simultaneously, and viruses were harvested 48h post transfection, filtered with 0.22 μm filters and store in aliquots at −80°C.The sequence of each gRNA used is reported in Table S2.

### RNAseq and analysis

Total RNA was extracted as described above. Sample for TruSeq stranded ribo-depleted paired-end total RNAseq at 40-50 million reads were submitted at the Epigenomic Core at Well Cornell Medical College (WCMC). Samples for paired-end poly-A enriched RNAseq at 20-30 million reads were submitted to the Memorial Sloan Kettering Cancer Center (MSKCC) Genomic Core. Quality control of sequenced reads was performed by FastQC. Adaptor-trimmed reads were mapped to the hg19 human genome using STAR ^52^. The htseq-count function of the HTSeq Python package ^53^ was used to count uniquely aligned reads at all exons of a gene. The count values were transformed to reads per kilobase per million (RPKM) to make them comparable across replicates. A threshold of 1 RPKM was used to consider a gene to be present in a sample and genes that were present in at least one sample were used for subsequent analyses. Differential gene expression across time-points was computed using DESeq2 ^54^. Variance stabilizing transformation (VST) of RNAseq counts was used for the Principal Component Analysis (PCA) Plots and for heatmaps of gene expression. For downstream analysis of trends of gene expression, transcripts were first grouped into “monotonically upregulated” and “monotonically downregulated” based on the characteristics of their expression from day25 to day100 and further categorized in *strict*: all the transitions satisfy the statistical significance criteria and *relaxed:* day25 *vs.* day100 transition satisfy the significance criteria and intermediate transitions may not. For all comparisons a significance threshold of FDR ≤ 5% was used. Monotonically upregulated (strict): (d50*vs.*d25: FDR ≤ 5%) AND (d100*vs.*d25: FDR ≤ 5%) AND (d100*vs.*d50: FDR ≤ 5%) AND (d50*vs.*d25: logFC > 0) AND (d75*vs.*d50: logFC > 0) AND (d100*vs.*d25 logFC > d50*vs.*d25 logFC). Monotonically downregulated (strict): (d50*vs.*d25: FDR ≤ 5%) AND (d100*vs.*d25: FDR ≤ 5%) AND (d100*vs.*d50: FDR ≤ 5%) AND (d50*vs.*d25: logFC < 0) AND (d75*vs.*d50: logFC < 0) AND (d100*vs.*d25 logFC < d50*vs.*d25 logFC). Monotonically upregulated (relaxed): (d100*vs.*d25: FDR ≤ 5%) AND (d50*vs.*d25: logFC > 0) AND ((d100*vs.*d25: logFC >= d50*vs.*d25: logFC) OR (d75*vs.*d50: logFC > 0)). Monotonically downregulated (relaxed): (d100*vs.*d25: FDR ≤ 5%) AND (d50*vs.*d25: logFC < 0) AND ((d100*vs.*d25: logFC <= d50*vs.*d25: logFC) OR (d75*vs.*d50: logFC < 0)). GSEA ^55^ was performed on day50 vs. day25 and day100 vs. day50 pairwise comparisons to test enrichment in KEGG pathways or gene sets from MSigDB using the following parameters: FDR ≤ 5%, minimum gene-set size=15, maximum gene-set size=500, number of permutations = 1000. GO analysis was performed using DAVID. Single-cell RNAseq analysis for mouse cortical development in Fig.3f,g derived from the published dataset by Di Bella et al.^33^. Data was processed using the same pipeline as in the original publication and developmental trajectories were inferred using URD algorithm^56^.

### ATACseq and analysis

ATACseq libraries were prepared at the Epigenetic Innovation Lab at MSKCC starting from ∼ 50,000 live adherent cells plated on 96-wells. Size-selected libraries were submitted to the MSKCC Genomic core for paired-end sequencing at 40-60 million reads. Quality control of sequenced reads was performed by FastQC and adaptor filtration was performed by Trimmomatic version 0.36. The filtered reads were aligned to the hg19 reference genome. Macs2 ^57^ was used for removing duplicate reads and calling peaks. Differentially accessible peaks in the atlas were called by DESeq2 ^54^. To define dynamic trends of chromatin accessibility during neuronal maturation as shown in Fig. 3G, agglomerative hierarchical clustering using Ward’s methods Ward’s linkage method was done on the union of differentially accessible peaks in pairwise comparisons between d25, d50, d75 and d100 samples. HOMER findMotifsGenome.pl ^58^ was used to investigate the motif enrichment in pairwise comparisons and unbiasedly clustered groups of peaks. Motif enrichment was also assessed by Kolmogorov-Smirnov and hypergeometric tests as previously described ^59^. ATAC-seq peaks in the atlas were associated with TF motifs in the an updated CIS-BP database ^60, 61^ using FIMO ^62^ of MEME suite ^63^. Hypergeometric test was used to compare the proportion of peaks containing a transcription factor motif in each group (foreground ratio) with that in the entire atlas (background ratio). Odds ratio represents the normalized enrichment of peaks associated with transcription factor motifs in the group compared to the background (foreground ratio/background ratio).

### Cut&Run and analysis

Cut&Run was performed from 50,000 cells per condition as previously described in ^64^ using the following antibodies: rabbit anti-H3K4me3 (aab8580, abcam); rabbit anti-H3K9me3 (ab8898, abcam); rabbit anti-H3K27me3 (9733, Cell Signaling Technologies); rabbit anti-H3K27ac (309034, Active Motif), normal rabbit IgG (2729, Cell Signaling Technologies). In brief, cells were harvested and bound to concanavalin A-coated magnetic beads after an 8min incubation at RT on a rotator. Cell membranes were permeabilized with digitonin and the different antibodies were incubated overnight at 4°C on a rotator. Beads were washed and incubated with pA-MN. Ca^2+^-induced digestion occurred on ice for 30min and stopped by chelation. DNA was finally isolated using an extraction method with phenol and chloroform as described in ^64^.

Sequencing reads were trimmed and filtered for quality and adapter content using version 0.4.5 of TrimGalore (https://www.bioinformatics.babraham.ac.uk/projects/trim_galore) and running version 1.15 of cutadapt and version 0.11.5 of FastQC. Reads were aligned to human assembly hg19 with version 2.3.4.1 of bowtie2 (http://bowtie-bio.sourceforge.net/bowtie2/index.shtml) and MarkDuplicates of Picard Tools version 2.16.0 was used for deduplication. Enriched regions were discovered using MACS2 with a p-value setting of 0.001 and a matched IgG as the control. The BEDTools suite (http://bedtools.readthedocs.io) was used to create normalized read density profiles. A global peak atlas was created by first removing blacklisted regions (https://www.encodeproject.org/annotations/ENCSR636HFF) then merging all peaks within 500 bp and counting reads with version 1.6.1 of featureCounts (http://subread.sourceforge.net). Reads were normalized by sequencing depth (to 10 million mapped fragments) and DESeq2 was used to calculate differential enrichment for all pairwise contrasts. Clustering was performed on the superset of differential peaks using k-means clustering by increasing k until redundant clusters arose. Gene annotations were created by assigning all intragenic peaks to that gene, and otherwise using linear genomic distance to transcription start site. The annotations in each cluster were used to intersect with the RNA-seq time series by plotting the average expression z-score of all peak-associated genes which are differentially expressed across any stage. Motif signatures and enriched pathways were obtained using Homer v4.11 (http://homer.ucsd.edu).

## Statistical analysis

Statistics were performed in PRISM (GraphPad) and R software. Data are represented as arithmetical means +/- standard error of the mean (s.e.m.) unless otherwise indicated.

---

**Extended Data Fig.1:**
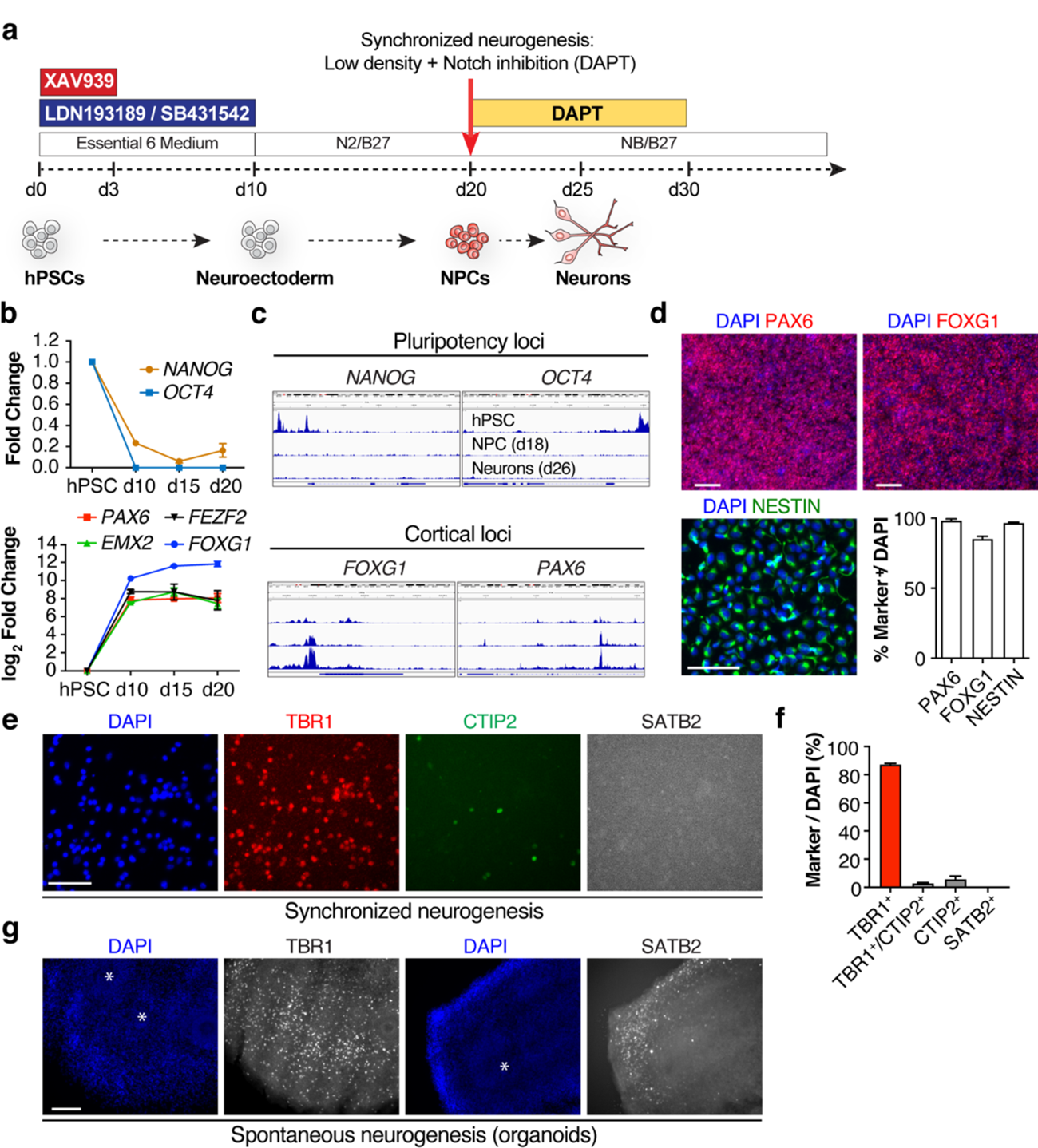
Generation of cortical neurons from hPSCs. **a**, Schematics of differentiation protocol based on dual-SMAD and WNT inhibition. Top panel indicate differentiation days, basal media, and small molecules treatments. Bottom panel indicate cell stages/types found at transition points. The red arrow indicates cell-passaging at low density in presence of notch pathway inhibitor DAPT. **b**, Expression of pluripotency (top) and cortical (bottom) specific markers by qRT-PCR throughout the differentiation (n = 3 independent experiments). **c**, Representative Genome browser traces of ATACseq peaks at hPSC, NPC and neuron stages in Pluripotency (*NANOG, OCT4*) and cortical (*PAX6, FOXG1*) loci (n = 2 independent experiments). **d**, Representative images and quantification of the fraction of cells expressing PAX6, FOXG1 and NESTIN cortical NPC markers at d20 of differentiation (n = 2 independent experiments). **e-f**, Representative images (e) and quantification (f) of the fraction of neurons generated through synchronized neurogenesis and expressing TBR1, CTIP2 and SATB2 cortical neuron markers. **g**, Representative images of neurons generated through spontaneous neurogenesis (cortical organoids) and stained with antibodies against cortical neurons markers (n = 2 independent experiments). * Marks rosette structures of neural precursor cells. Histograms depict mean ± s.e.m. Scale bars are 50 μm (e); 100 μm (d) and 200 μm (g).

**Extended Data Fig.2:**
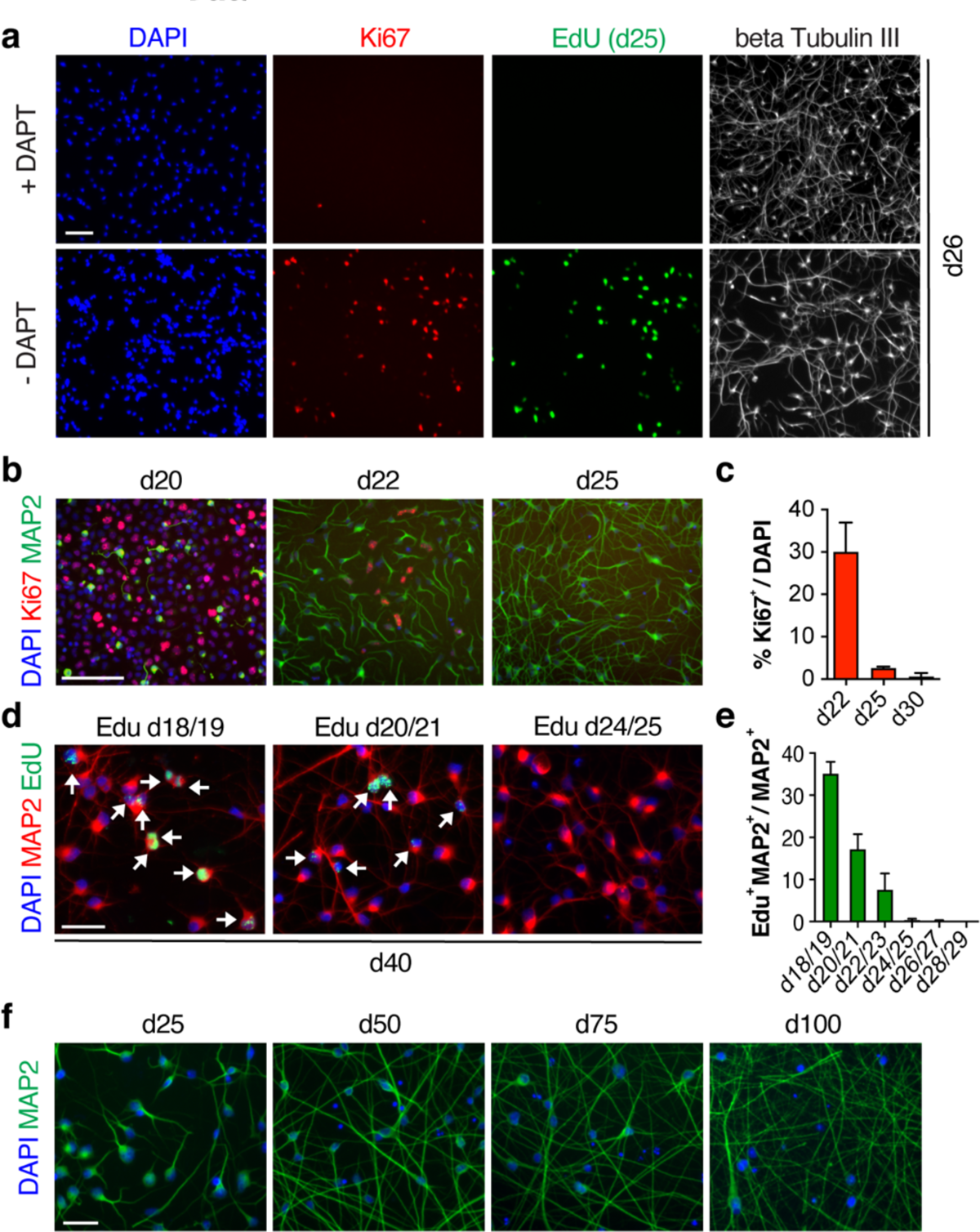
A novel platform for the synchronized neurogenesis of cortical neurons from hPSCs. **a**, Cell passaging at low density and DAPT treatment rapidly deplete the pool of progenitor cells. Cells were culture in presence or absence of DAPT from d20, pulse labelled with EdU for 24h at d25 and analyzed at d26 by immunostaining for EdU, KI67 and MAP2. **b-c**, Representative images (b) and quantification (c) of percentage of MKI67^+^ NPC and MAP2^+^ neurons after the induction of synchronized neurogenesis at d20 (n = 2 independent experiments). **d-e**, Representative images (d) and quantification (e) of the fraction of d40 MAP2 neurons that were labelled by EdU pulses of progenitor cells at the indicated days (n = 2 independent experiments). **f**, Synchronized cortical neurons can be maintained at high viability in long-term cultures. Representative images of cortical neurons stained with antibody against MAP2 at day25, 50, 75 and 100 of differentiation. Histograms depict mean ± s.e.m. Scale bars are 100 μm (a,b); 50 μm (d,f).

**Extended Data Fig.3:**
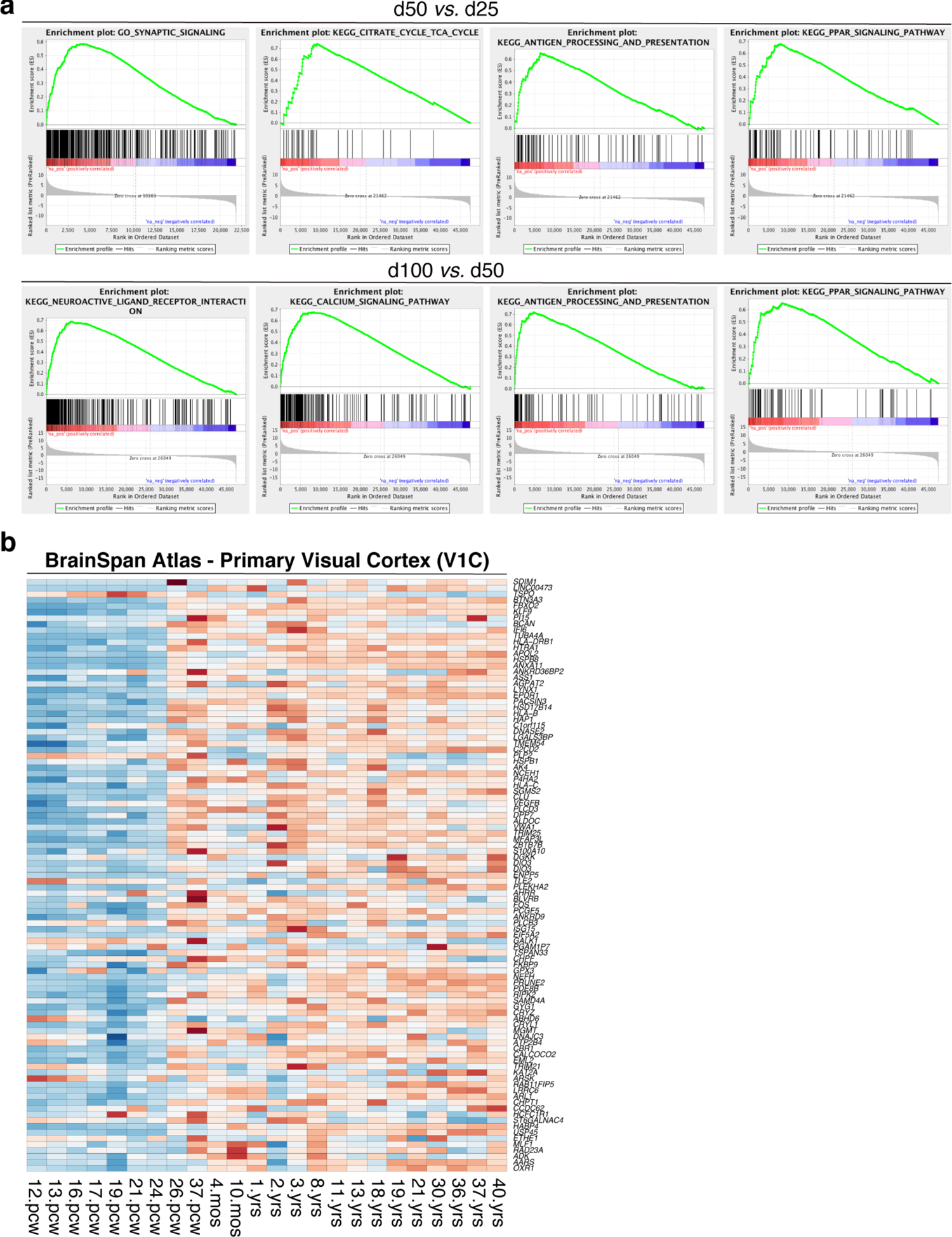
Gene ontology and BrainSpan comparison for maturation dependent transcripts. **a**, GSEA plots for some of the GO terms from RNAseq studies (n = 3 independent experiments) related that positively correlate with neuronal maturation in d50 *vs.* d25 and d100 *vs*. d50 pairwise comparisons. **b,** Heatmap for the normalized temporal expression of the corresponding monotonically upregulated transcripts in the BrainSpan atlas of the developing human brain (primary visual cortex) shown in Fig. 3c.

**Extended Data Fig.4:**
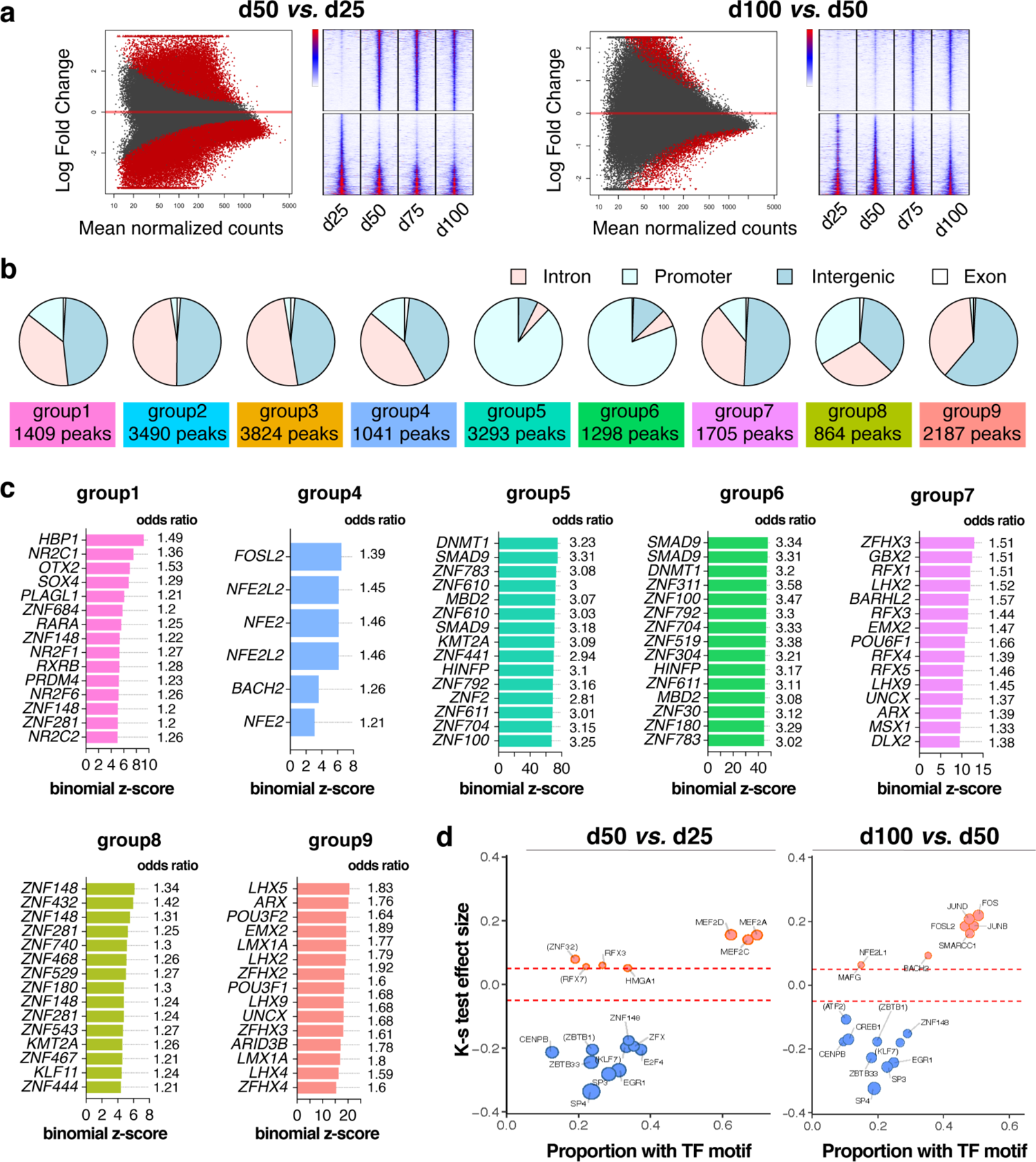
Motif analysis for unbiassed ATACseq clusters and pairwise comparisons of chromatin accessibility during maturation. **a**, Pie charts of ATACseq peaks mapped to gene promoters, introns, exons, and intergenic genomic regions for each of the cluster in Fig. 3f. **b**, Top transcription factor motifs enriched in the indicated groups of ATACseq peaks. Odds ratio indicates the normalized enrichment of transcription factor motifs in the cluster compared to the background. **c,** MA (left) and tornado plots (right) for differential accessible ATACseq peaks in d25 *vs.* d50 and d50 *vs*. d100 pairwise comparisons. **d,** Top transcription factor motifs enriched in differentially accessible ATACseq peaks in d50 *vs*. d25 and d100 *vs*. d50 pairwise comparisons. (a-d) n = 2 independent experiments.

**Extended Data Fig.5:**
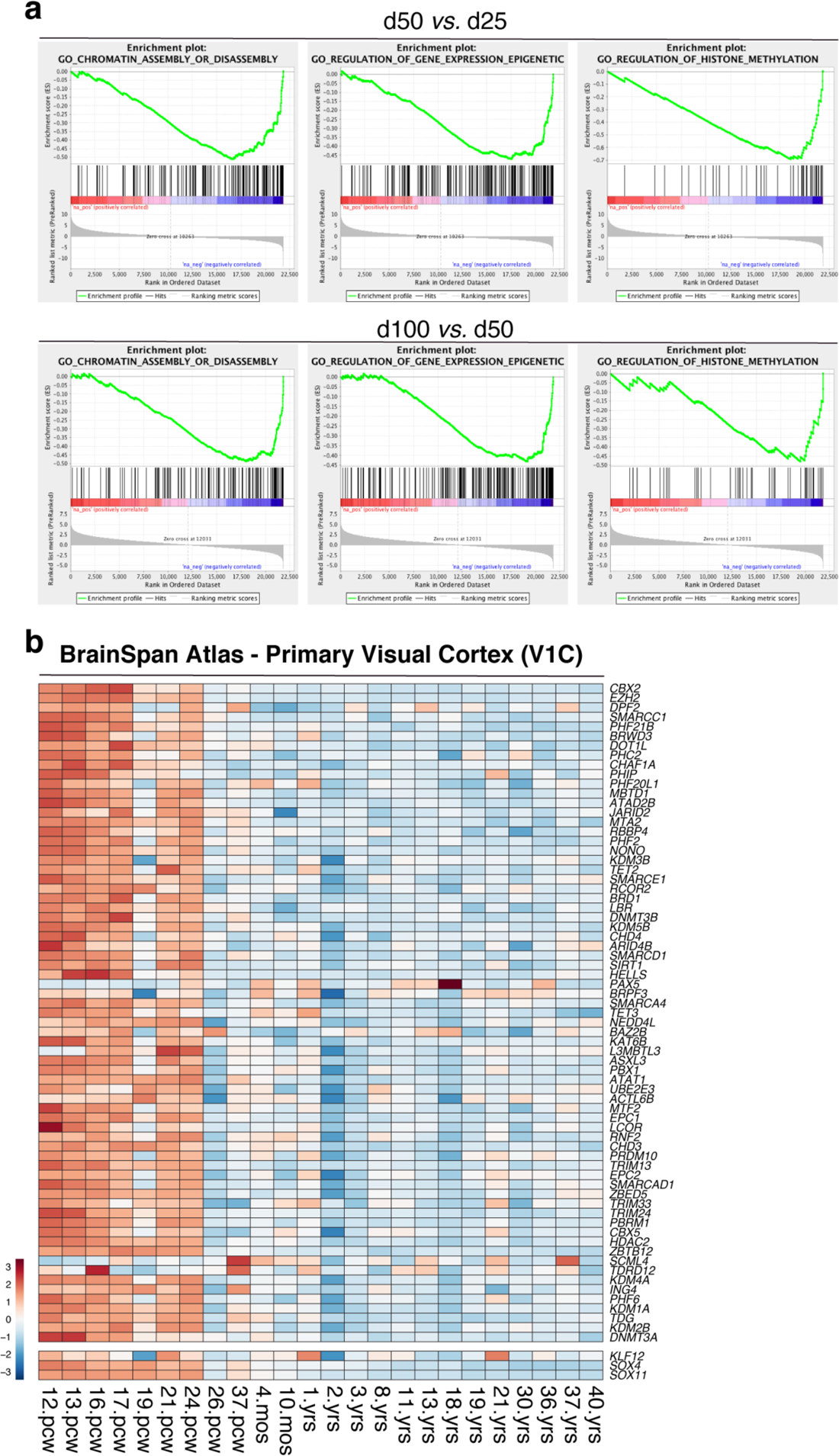
Chromatin regulators are progressively downregulated during neuronal maturation. **a**, GSEA plots for some of the GO terms from RNAseq studies (n = 3 independent experiments) related to chromatin remodeling in d50 *vs.* d25 and d100 *vs*. d50 pairwise comparisons. **b,** Heatmap for the normalized temporal expression of the corresponding monotonically downregulated chromatin regulators in the BrainSpan atlas of the developing human brain (primary visual cortex) shown in Fig. 4B.

**Extended Data Fig.6:**
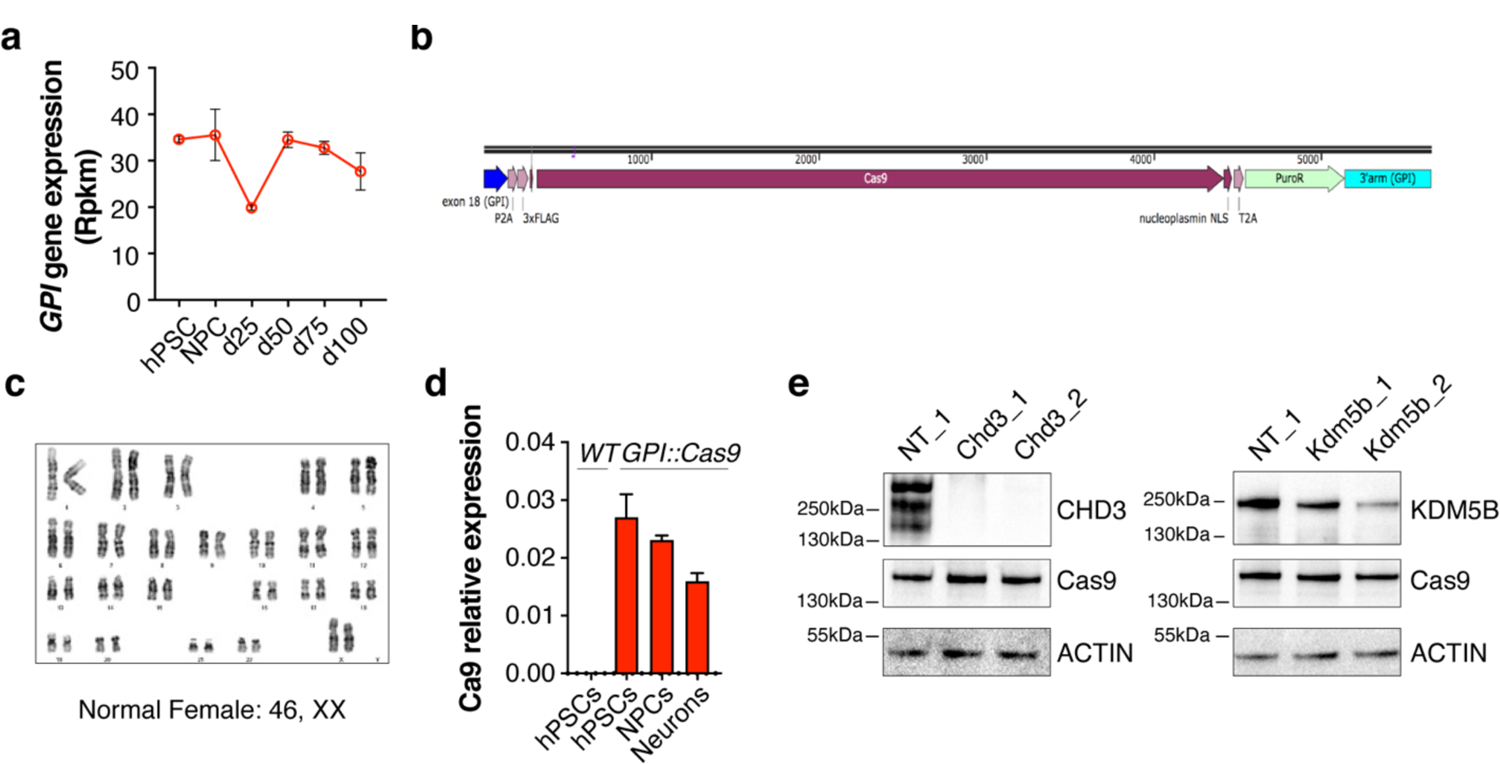
Experimental strategy for gene knock-out in hPSCs-derived neurons. **a**, Expression of *GPI* gene throughout the differentiation (n = 3 independent experiments). **b,** Schematic for the targeting construct used for the generation of the GPI::Cas9 knock-in hPSCs line. Cas9 is linked to the GPI gene via 2A self-cleaving peptide sequence. **c,** Karyotypic analysis of the GPI::Cas9 hPSCs clonal cell line used for the study. **d,** Expression of Cas9 mRNA in the GPI::Cas9 line at hPSC, NPC and neuron stages compared to wild type hPSC (n = 2 independent experiments). **e,** Representative Western Blot analysis for CRISPR/Cas9-based gene KO for CHD3 and KDM5B in neurons using the same strategy shown in Fig. 3c. Cas9 expressing neurons at d25 were infected with lentiviral vectors encoding non-targeting and gene-specific gRNAs and analyzed at day35 of differentiation. Histograms depict mean ± s.e.m.

**Extended Data Fig.7:**
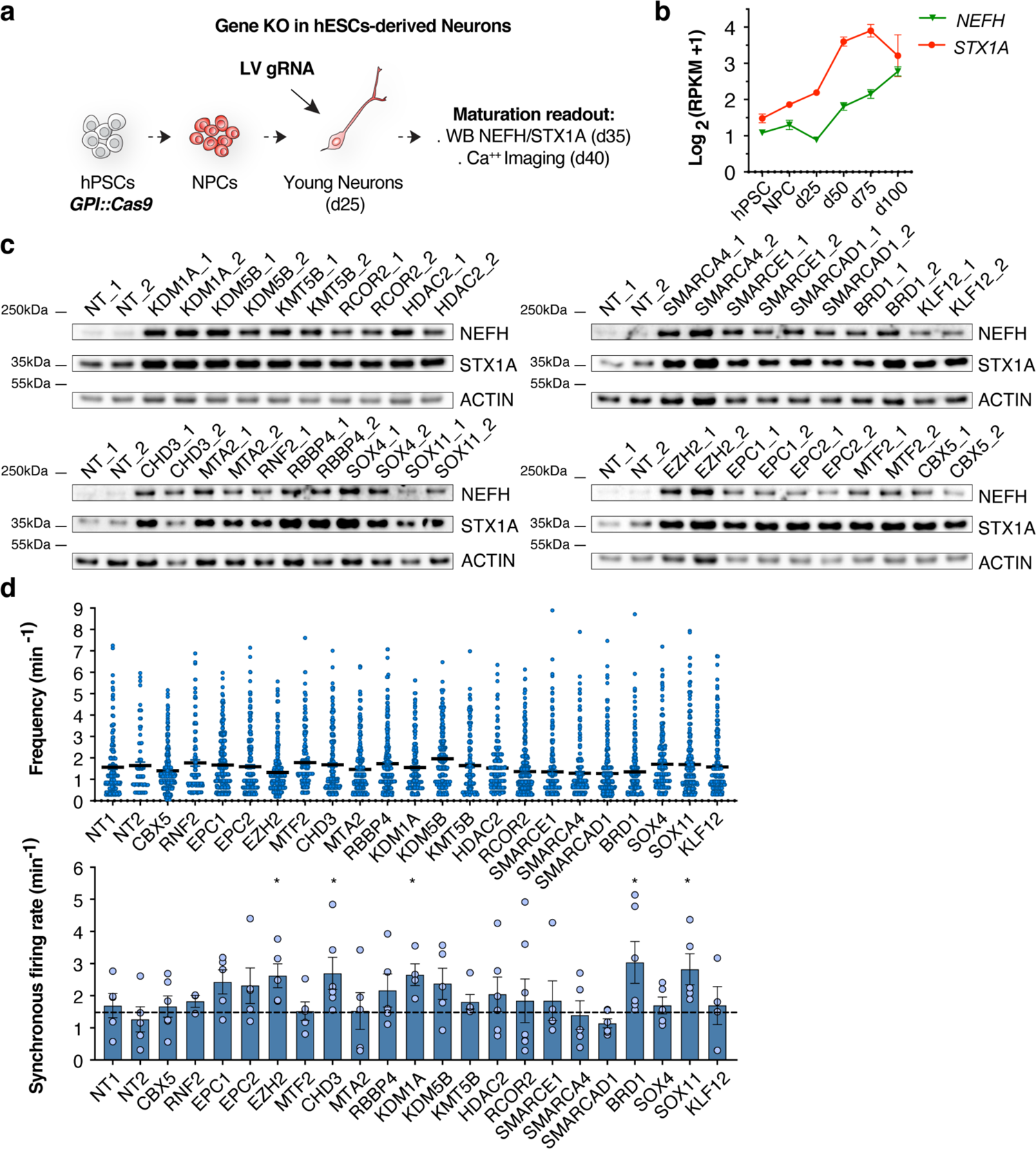
Loss-of-function of epigenetic factors in neurons induce preconscious maturation. **a**, Schematic of experimental paradigm for gene KO in hPSCs-derived neurons. **b,** Gene expression for NEFH and STX1A maturation markers throughout the differentiation (n = 3 independent experiments). **c,** Representative Western Blots for the loss-of-function genetic screen of chromatin regulators. **d,** Frequency of spontaneous Ca^2+^ spikes of individual neurons (top) and synchronicity rate of spontaneous network activity (bottom) in loss-of-function experiments. Dots represent individual neurons and FOW respectively (n = 3-6 FOW from 2 independent experiments). Histograms and lines depict mean ± s.e.m. Kruskal-Wallis test; *p < 0.05.

**Extended Data Fig.8:**
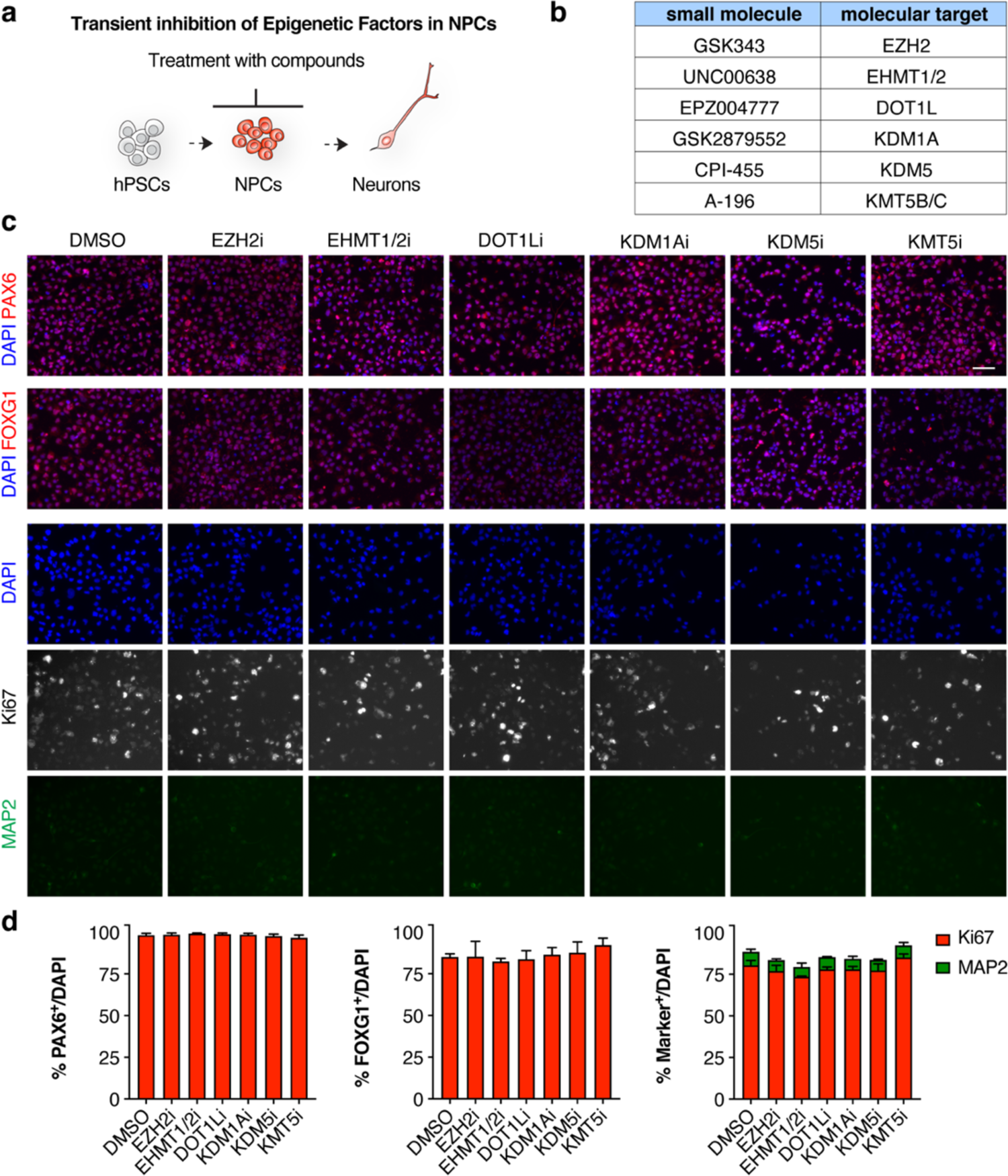
Transient inhibition of epigenetic factors in NPC does not alter cortical patterning and neurogenesis. **a**, Schematic of experimental paradigm for transient inhibition of chromatin regulators at progenitor cell stage. NPC were treated with small molecule from d12 to d20. **b,** Small molecule compounds used in the study and corresponding intracellular targets. **c,** Representative images of d20 NPCs treated with small molecule before the induction of synchronized neurogenesis and stained with antibodies against cortical markers PAX6 and FOXG1, the proliferation marker Ki67 and the neuron marker MAP2. **d,** Quantification of the fraction of cells expressing each marker in treated *vs.* control conditions (n = 2 independent experiments). Histograms depict mean ± s.e.m. Scale bars are 50 μm.

**Extended Data Fig.9:**
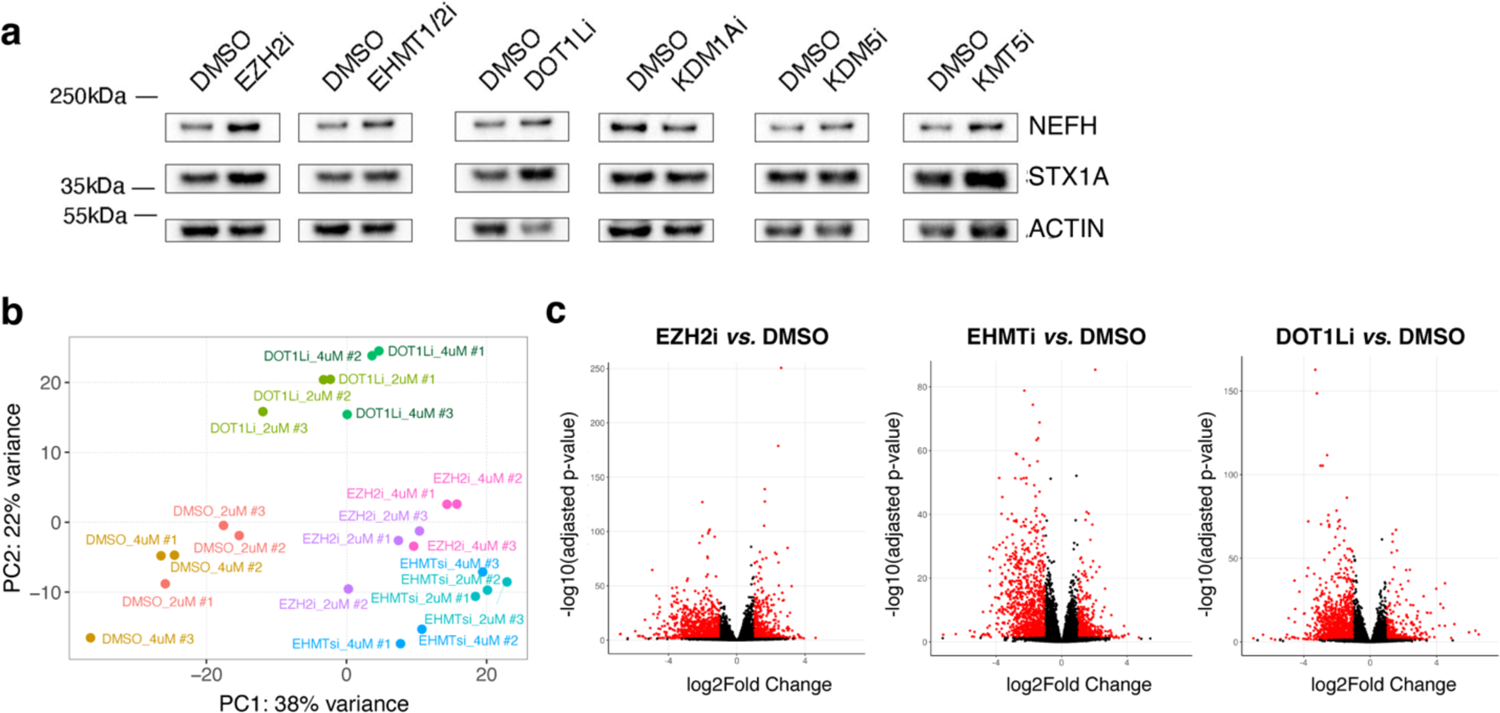
A small molecule mini screen identify EZH2, EHMT1/2 and DOT1L inhibition in NPC as a maturation driver in neurons. **a**, Representative Western Blots for the expression of NEFH and STX1A maturation markers in the transient inhibition of epigenetic factors in NPC experiments. NPC were treated with small molecule from d12 to d20 and neurons derived from each condition were analyzed at d35**. b,** PCA plot for RNAseq dataset show samples distribution according to the pharmacological treatments (n = 3 independent experiments). **c,** Volcano plot for the indicated pairwise comparisons from RNAseq studies (treatment with 4μM of inhibitors). Red dots represent differentially expressed significant transcript (FDR 0.05) that show Fold Change >=2.

**Extended Data Fig.10:**
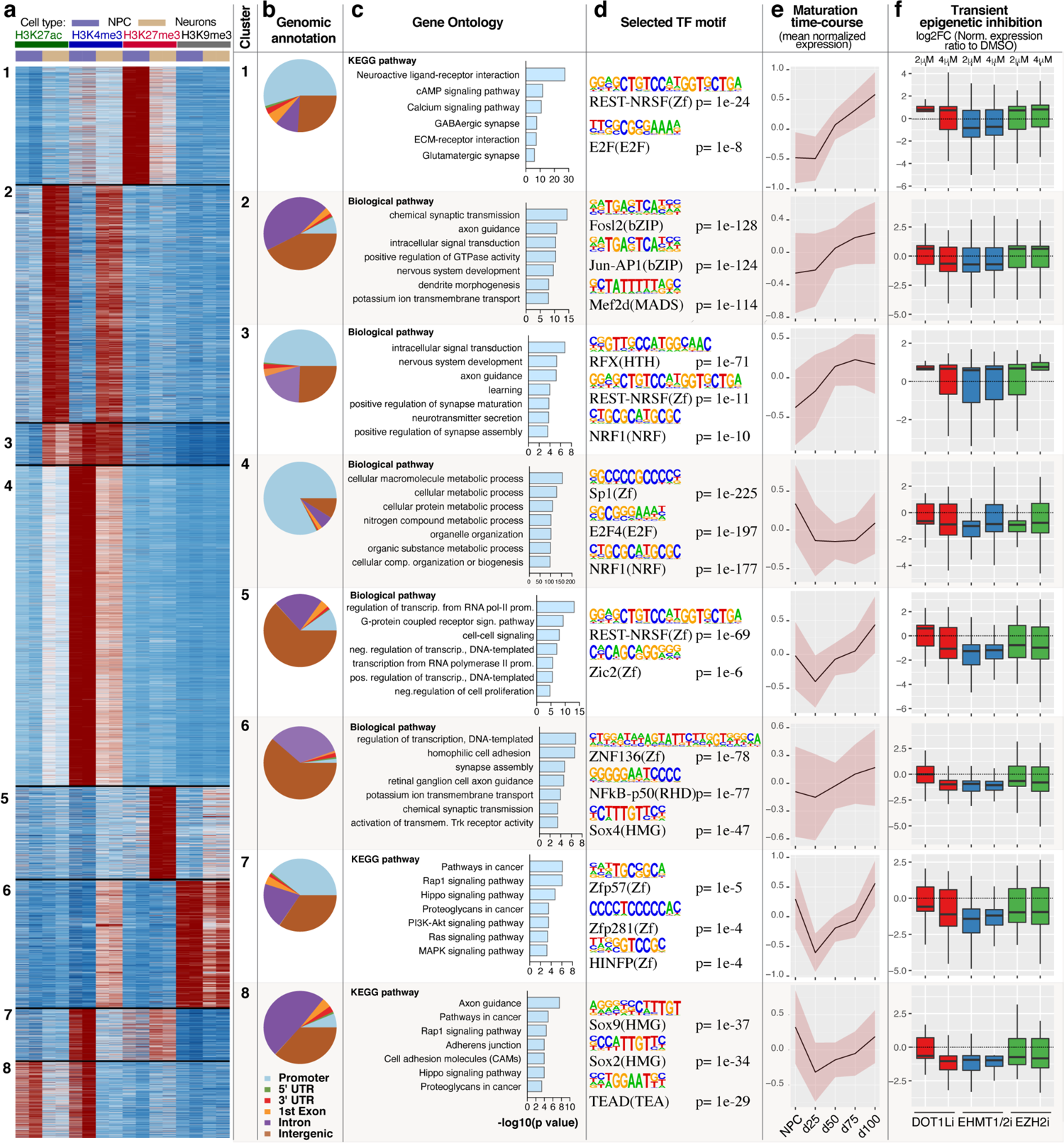
Patterns of histone post translational modifications drive the maturation of hPSC-derived neurons. **a**, Unsupervised clustering of CUT&RUN peaks with differential density in H3K27ac, H3K4me4, H3K27me3 and H3K9me3 signal among NPC and neurons (n = 2 replicates/condition). **b,** Pie charts of CUT&RUN peaks mapped to gene promoters, introns, exons, and intergenic genomic regions for each of the cluster. **c,** GO for genes linked at each cluster. **d,** Top selected statistically significant enriched transcription factor motifs at peaks in each cluster. **e,** Mean normalized expression (z-transform) of differentially expressed genes during the maturation time course intersected with genes linked to each CUT&RUN cluster. **f,** Expression of differentially expressed transcripts from (e) in neurons derived from NPC treated with the indicated inhibitors respect to DMSO controls. Pink area in (e) is S.E.M. and whiskers in (e) depict 1.5*interquartile range beyond the 25^th^ and 75^th^ percentiles.

**Table S1.** Electrophysiological properties of maturing hPSC-derived cortical neurons. Quantification of the electrophysiological properties of hPSC-derived neurons at day 25, 50, 75, 100 of differentiation. Results are displayed as mean ± s.e.m. RMP, Resting membrane potential; IR, Input resistance, APT, Action potential threshold; APA, Action potential amplitude; APD, Action potential duration at its half amplitude; AHP, After hyperpolarization amplitude; Cm, Membrane capacitance; Rm, Membrane resistance.

|  | day 25 | day 50 | day 75 | day 100 |
| --- | --- | --- | --- | --- |
| RMP (mV) | -36.7±2.0 | -50.2±1.2 | -52.3±1.1 | -51.6±1.4 |
| IR (MΩ) | 1,601.5±100.9 | 731.4±53.3 | 646.0±48.6 | 518.5±65.9 |
| Rheobase (pA) | 98.8±8.5 | 32.7±6.5 | 36.0±4.7 | 29.7±4.5 |
| APT (mV) | -8.8±2.4 | -31.5±1.2 | -30.1±1.0 | -34.5±1.0 |
| APA (mV) | 25.1±1.3 | 48.2±3.1 | 53.1±1.8 | 59.1±2.3 |
| APD (ms) | 23.9±2.1 | 14.1±1.4 | 10.0±1.0 | 9.5±0.7 |
| AHP (ms) | 12.0±1.2 | 6.0±0.7 | 4.0±0.3 | 3.5±0.2 |
| Rise time (ms) | 5.5±0.5 | 2.5±0.3 | 1.7±0.2 | 1.6±0.2 |
| Decay time (ms) | 9.6±0.9 | 7.3±1.0 | 4.3±0.5 | 4.1±0.3 |
| Rise slope (mV/ms) | 4.4±0.8 | 29.7±4.3 | 39.8±4.0 | 42.4±4.3 |
| Decay slope (mV/ms) | -2.9±0.7 | -9.9±1.4 | -14.2±1.3 | -13.7±1.1 |
| Cm (pF) | 21.2±1.6 | 28.3±1.9 | 35.2±2.0 | 31.6±1.9 |
| Rm (MΩ) | 505.2±61.6 | 192.0±20.0 | 144.8±8.7 | 143.6±13.5 |
| Access resistance (MΩ) | 39.8±2.6 | 42.9±2.3 | 36.5±1.9 | 39.6±1.7 |
| n | 25 | 33 | 43 | 29 |

**Table S2.**
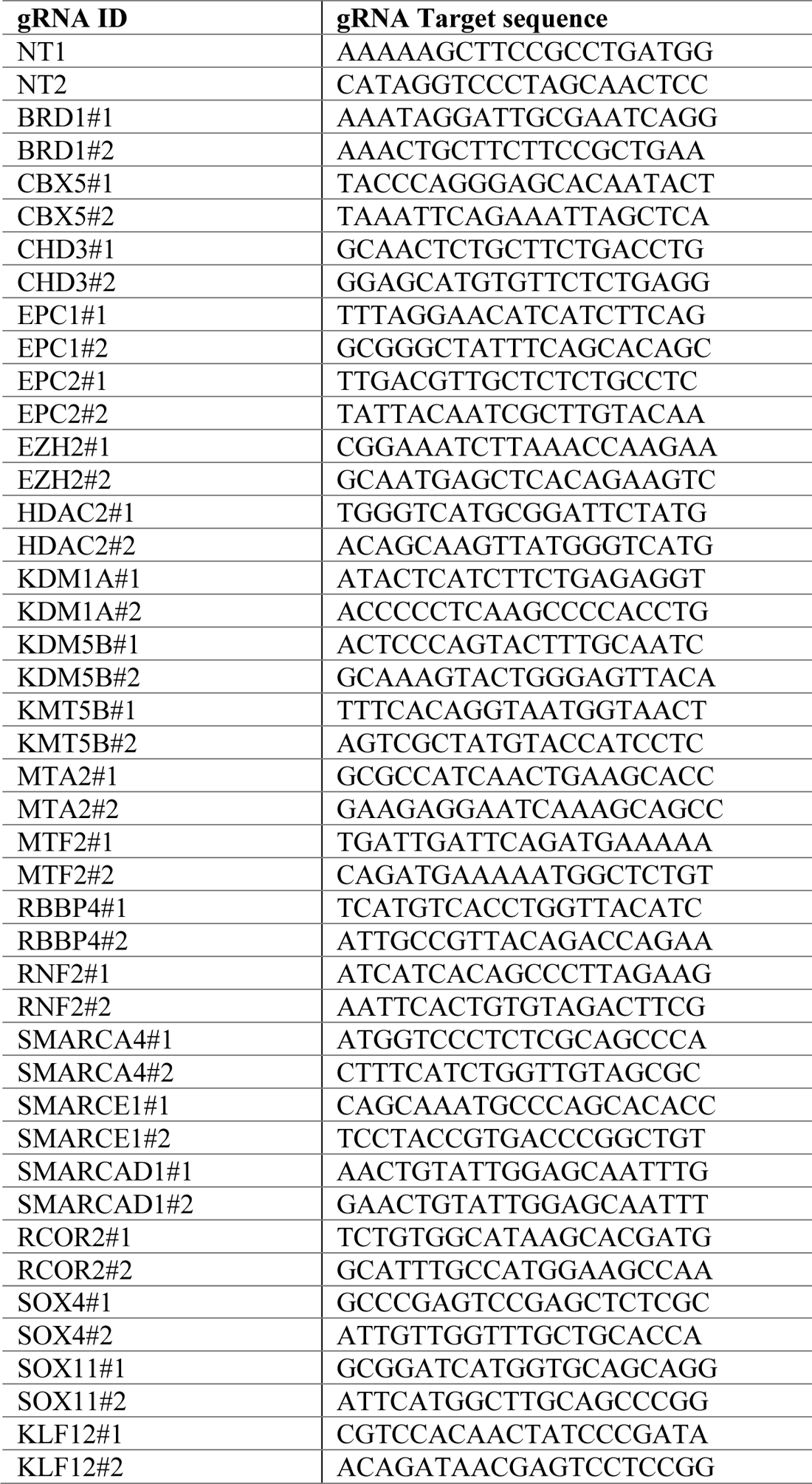
**List of gRNA sequences used for gene KO in hPSC-derived neurons.**

## Supplementary Information

**Movie S1.** Representative Ca^2+^ imaging for hPSC-derived cortical neurons at d40 of differentiation.

**Movie S2.** Representative Ca^2+^ imaging for hPSC-derived cortical neurons at d70 of differentiation.

**Movie S3.** Representative Ca^2+^ imaging for hPSC-derived cortical neurons at d40 derived from control (DMSO) treated NPC

**Movie S4.** Representative Ca^2+^ imaging for hPSC-derived cortical neurons at d40 derived from NPC treated EZH2 inhibitor

**Movie S5.** Representative Ca^2+^ imaging for hPSC-derived cortical neurons at d40 derived from NPC treated EHMT1/2 inhibitor

**Movie S6.** Representative Ca^2+^ imaging for hPSC-derived cortical neurons at d40 derived from NPC treated DOT1L inhibitor

## References

1. Toma, K., Wang, T. C. & Hanashima, C. Encoding and decoding time in neural development. Dev Growth Differ 58, 59–72, doi:10.1111/dgd.12257 (2016).

2. Ebisuya, M. & Briscoe, J. What does time mean in development? Development 145, doi:10.1242/dev.164368 (2018).

3. Zhu, Y. et al. Spatiotemporal transcriptomic divergence across human and macaque brain development. Science 362, doi:10.1126/science.aat8077 (2018).

4. Marchetto, M. C. et al. Species-specific maturation profiles of human, chimpanzee and bonobo neural cells. Elife 8, doi:10.7554/eLife.37527 (2019).

5. Barry, C. et al. Species-specific developmental timing is maintained by pluripotent stem cells ex utero. Dev Biol 423, 101–110, doi:10.1016/j.ydbio.2017.02.002 (2017).

6. Gaspard, N. et al. in Nature Vol. 455 351–357 (2008).

7. Espuny-Camacho, I. et al. in Neuron Vol. 77 440–456 (2013).

8. Shi, Y., Kirwan, P., Smith, J., Robinson, H. P. C. & Livesey, F. J. in Nature Neuroscience Vol. 15 477–486, S471 (2012).

9. Otani, T., Marchetto, M. C., Gage, F. H., Simons, B. D. & Livesey, F. J. in Cell Stem Cell Vol. 18 467–480 (2016).

10. Linaro, D. et al. Xenotransplanted Human Cortical Neurons Reveal Species-Specific Development and Functional Integration into Mouse Visual Circuits. Neuron 104, 972–986 e976, doi:10.1016/j.neuron.2019.10.002 (2019).

11. Rayon, T. et al. Species-specific pace of development is associated with differences in protein stability. Science 369, doi:10.1126/science.aba7667 (2020).

12. Matsuda, M. et al. Species-specific segmentation clock periods are due to differential biochemical reaction speeds. Science 369, 1450-+, doi:10.1126/science.aba7668 (2020).

13. Cardoso-Moreira, M. et al. Gene expression across mammalian organ development. Nature 571, 505–509, doi:10.1038/s41586-019-1338-5 (2019).

14. Barry, C. et al. Automated minute scale RNA-seq of pluripotent stem cell differentiation reveals early divergence of human and mouse gene expression kinetics. PLoS Comput Biol 15, e1007543, doi:10.1371/journal.pcbi.1007543 (2019).

15. Sousa, A. M. M., Meyer, K. A., Santpere, G., Gulden, F. O. & Sestan, N. Evolution of the Human Nervous System Function, Structure, and Development. Cell 170, 226–247, doi:10.1016/j.cell.2017.06.036 (2017).

16. Silbereis, J. C., Pochareddy, S., Zhu, Y., Li, M. & Sestan, N. The Cellular and Molecular Landscapes of the Developing Human Central Nervous System. Neuron 89, 248–268, doi:10.1016/j.neuron.2015.12.008 (2016).

17. Kriks, S. et al. Dopamine neurons derived from human ES cells efficiently engraft in animal models of Parkinson’s disease. Nature 480, 547–551, doi:10.1038/nature10648 (2011).

18. Chambers, S. M. et al. in Nat Biotechnol Vol. 30 715–720 (2012).

19. Maroof, A. M. et al. in Cell Stem Cell Vol. 12 559–572 (2013).

20. Nicholas, C. R. et al. in Cell Stem Cell Vol. 12 573–586 (2013).

21. Ullian, E. M., Sapperstein, S. K., Christopherson, K. S. & Barres, B. A. Control of synapse number by glia. Science 291, 657–661, doi:10.1126/science.291.5504.657 (2001).

22. Tchieu, J. et al. NFIA is a gliogenic switch enabling rapid derivation of functional human astrocytes from pluripotent stem cells. Nat Biotechnol 37, 267–275, doi:10.1038/s41587-019-0035-0 (2019).

23. West, A. E. & Greenberg, M. E. Neuronal activity-regulated gene transcription in synapse development and cognitive function. Cold Spring Harb Perspect Biol 3, doi:10.1101/cshperspect.a005744 (2011).

24. Huang, E. J. & Reichardt, L. F. Neurotrophins: roles in neuronal development and function. Annu Rev Neurosci 24, 677–736, doi:10.1146/annurev.neuro.24.1.677 (2001).

25. Qi, Y. et al. Combined small-molecule inhibition accelerates the derivation of functional cortical neurons from human pluripotent stem cells. Nat Biotechnol 35, 154–163, doi:10.1038/nbt.3777 (2017).

26. Tornero, D. et al. Human induced pluripotent stem cell-derived cortical neurons integrate in stroke-injured cortex and improve functional recovery. Brain 136, 3561–3577, doi:10.1093/brain/awt278 (2013).

27. Marin, O. Developmental timing and critical windows for the treatment of psychiatric disorders. Nat Med 22, 1229–1238, doi:10.1038/nm.4225 (2016).

28. Shen, Q. et al. The timing of cortical neurogenesis is encoded within lineages of individual progenitor cells. Nat Neurosci 9, 743–751, doi:10.1038/nn1694 (2006).

29. Ben-Ari, Y. in Nature Reviews Neuroscience Vol. 3 728–739 (2002).

30. Bar-Shira, O., Maor, R. & Chechik, G. Gene Expression Switching of Receptor Subunits in Human Brain Development. PLoS Comput Biol 11, e1004559, doi:10.1371/journal.pcbi.1004559 (2015).

31. Zheng, X. et al. Metabolic reprogramming during neuronal differentiation from aerobic glycolysis to neuronal oxidative phosphorylation. Elife 5, doi:10.7554/eLife.13374 (2016).

32. Kang, H. J. et al. Spatio-temporal transcriptome of the human brain. Nature 478, 483–489, doi:10.1038/nature10523 (2011).

33. Di Bella, D. J. et al. Molecular logic of cellular diversification in the mouse cerebral cortex. Nature 595, 554–559, doi:10.1038/s41586-021-03670-5 (2021).

34. Flavell, S. W. et al. Activity-dependent regulation of MEF2 transcription factors suppresses excitatory synapse number. Science 311, 1008–1012, doi:10.1126/science.1122511 (2006).

35. Pajares, M. et al. Transcription factor NFE2L2/NRF2 is a regulator of macroautophagy genes. Autophagy 12, 1902–1916, doi:10.1080/15548627.2016.1208889 (2016).

36. Malik, A. N. et al. Genome-wide identification and characterization of functional neuronal activity-dependent enhancers. Nat Neurosci 17, 1330–1339, doi:10.1038/nn.3808 (2014).

37. Bernstein, B. E. et al. A bivalent chromatin structure marks key developmental genes in embryonic stem cells. Cell 125, 315–326, doi:10.1016/j.cell.2006.02.041 (2006).

38. Han, X. et al. Destabilizing LSD1 by Jade-2 promotes neurogenesis: an antibraking system in neural development. Mol Cell 55, 482–494, doi:10.1016/j.molcel.2014.06.006 (2014).

39. Potts, R. C. et al. CHD5, a brain-specific paralog of Mi2 chromatin remodeling enzymes, regulates expression of neuronal genes. PLoS One 6, e24515, doi:10.1371/journal.pone.0024515 (2011).

40. Charrier, C. et al. in Cell Vol. 149 923–935 (2012).

41. Fossati, M. et al. SRGAP2 and Its Human-Specific Paralog Co-Regulate the Development of Excitatory and Inhibitory Synapses. Neuron 91, 356–369, doi:10.1016/j.neuron.2016.06.013 (2016).

42. Fiddes, I. T. et al. Human-Specific NOTCH2NL Genes Affect Notch Signaling and Cortical Neurogenesis. Cell 173, 1356–1369 e1322, doi:10.1016/j.cell.2018.03.051 (2018).

43. Suzuki, I. K. et al. Human-Specific NOTCH2NL Genes Expand Cortical Neurogenesis through Delta/Notch Regulation. Cell 173, 1370–1384 e1316, doi:10.1016/j.cell.2018.03.067 (2018).

44. Chen, C., Lee, G. A., Pourmorady, A., Sock, E. & Donoghue, M. J. Orchestration of Neuronal Differentiation and Progenitor Pool Expansion in the Developing Cortex by SoxC Genes. J Neurosci 35, 10629–10642, doi:10.1523/JNEUROSCI.1663-15.2015 (2015).

45. Klingler, E. et al. Temporal controls over inter-areal cortical projection neuron fate diversity. Nature, doi:10.1038/s41586-021-04048-3 (2021).

46. Egan, C. M. et al. in Developmental Cell Vol. 26 223–236 (2013).

47. Xie, C. R. et al. Mutual regulation between CHD5 and EZH2 in hepatocellular carcinoma. Oncotarget 6, 40940–40952, doi:10.18632/oncotarget.5724 (2015).

48. Telley, L. et al. Temporal patterning of apical progenitors and their daughter neurons in the developing neocortex. Science 364, doi:10.1126/science.aav2522 (2019).

49. Pereira, J. D. et al. Ezh2, the histone methyltransferase of PRC2, regulates the balance between self-renewal and differentiation in the cerebral cortex. Proc Natl Acad Sci U S A 107, 15957–15962, doi:10.1073/pnas.1002530107 (2010).

50. Sun, Z. & Sudhof, T. C. A simple Ca(2+)-imaging approach to neural network analyses in cultured neurons. J Neurosci Methods 349, 109041, doi:10.1016/j.jneumeth.2020.109041 (2021).

51. Concordet, J. P. & Haeussler, M. CRISPOR: intuitive guide selection for CRISPR/Cas9 genome editing experiments and screens. Nucleic Acids Res 46, W242–W245, doi:10.1093/nar/gky354 (2018).

52. Dobin, A. et al. STAR: ultrafast universal RNA-seq aligner. Bioinformatics 29, 15–21, doi:10.1093/bioinformatics/bts635 (2013).

53. Anders, S., Pyl, P. T. & Huber, W. HTSeq--a Python framework to work with high-throughput sequencing data. Bioinformatics 31, 166–169, doi:10.1093/bioinformatics/btu638 (2015).

54. Love, M. I., Huber, W. & Anders, S. Moderated estimation of fold change and dispersion for RNA-seq data with DESeq2. Genome Biol 15, 550, doi:10.1186/s13059-014-0550-8 (2014).

55. Subramanian, A. et al. Gene set enrichment analysis: a knowledge-based approach for interpreting genome-wide expression profiles. Proc Natl Acad Sci U S A 102, 15545–15550, doi:10.1073/pnas.0506580102 (2005).

56. Farrell, J. A. et al. Single-cell reconstruction of developmental trajectories during zebrafish embryogenesis. Science 360, doi:10.1126/science.aar3131 (2018).

57. Zhang, Y. et al. Model-based analysis of ChIP-Seq (MACS). Genome Biol 9, R137, doi:10.1186/gb-2008-9-9-r137 (2008).

58. Heinz, S. et al. Simple combinations of lineage-determining transcription factors prime cis-regulatory elements required for macrophage and B cell identities. Mol Cell 38, 576–589, doi:10.1016/j.molcel.2010.05.004 (2010).

59. Lee, K. et al. FOXA2 Is Required for Enhancer Priming during Pancreatic Differentiation. Cell Rep 28, 382–393 e387, doi:10.1016/j.celrep.2019.06.034 (2019).

60. Weirauch, M. T. et al. Determination and inference of eukaryotic transcription factor sequence specificity. Cell 158, 1431–1443, doi:10.1016/j.cell.2014.08.009 (2014).

61. Lambert, S. A. et al. The Human Transcription Factors. Cell 175, 598–599, doi:10.1016/j.cell.2018.09.045 (2018).

62. Grant, C. E., Bailey, T. L. & Noble, W. S. FIMO: scanning for occurrences of a given motif. Bioinformatics 27, 1017–1018, doi:10.1093/bioinformatics/btr064 (2011).

63. Bailey, T. L. et al. MEME SUITE: tools for motif discovery and searching. Nucleic Acids Res 37, W202–208, doi:10.1093/nar/gkp335 (2009).

64. Skene, P. J., Henikoff, J. G. & Henikoff, S. Targeted in situ genome-wide profiling with high efficiency for low cell numbers. Nat Protoc 13, 1006–1019, doi:10.1038/nprot.2018.015 (2018).

